# Transgenerational polarity axis inheritance during *Ceratopteris* embryogenesis

**DOI:** 10.1101/2025.08.29.673061

**Authors:** Sjoerd Woudenberg, Andrew R.G. Plackett, Zhaodong Hao, Hidemasa Suzuki, Luis Alonso Baez, Cecilia Borassi, Thorsten Hamann, Minako Ueda, Jane A. Langdale, Joris Sprakel, Jasper van der Gucht, Dolf Weijers

## Abstract

For sexually reproducing organisms to pass on their genetic information, progeny must successfully establish. Various life history strategies have evolved, using either dispersal of large numbers of progeny or intensive nurturing of a few. Most plants use the former strategy, but ferns generate a single embryo in the exact same location as the mother, and it is unknown how progeny success is promoted, or how embryogenesis is adapted. By studying *Ceratopteris richardii* embryogenesis, we find that maternal tissues guide orientation of the early embryo body axis, thus aligning its root pole towards the homologous maternal rhizoids. We find that axis polarity inheritance is mediated by maternal tissue mechanical patterns, and thus identify a robust mechanism for progeny establishment as a nurturing strategy in plants.

## Main Text

Evolutionary success of species depends on the transgenerational propagation of variants and traits. A key question, therefore, is how organisms ensure establishment and survival of their offspring. Sexual reproduction is a crucial moment during an organism’s life cycle and reproductive strategies are deeply intertwined with different life histories (*1*). The investment in individual offspring is highly diverse among sexually reproducing eukaryotes, ranging from masses of spores in fungi (*2, 3*), thousands of eggs in amphibians and fishes (*4, 5*), or a handful of offspring in mammals and birds (*6, 7*). In non-motile plants, reproductive life histories are more constrained due to high fitness costs related to dispersal (*8, 9*). There are two dominant reproductive life history strategies: (1) in bryophytes, the fertilization product (embryo) immediately generates numerous somatic cells that collectively enter meiosis and disperse as spores; (2) in seed plants, embryogenesis occurs within the seed, which, following dispersal, will establish a progeny plant elsewhere. Seed plants generally produce large numbers of seeds to ensure successful establishment of some, and have evolved many adaptions for dispersal (*10–13*). In both cases, large numbers of offspring are produced in a “hail shot” strategy to deal with the uncertainty of progeny establishment, yet the function of the sporophyte (embryo) is dramatically different, and this difference may impact on the nurturing strategy chosen for the progeny.

Seed plants are sister to ferns, a group of seedless vascular plants whose life cycle combines a free-living bryophyte-like indeterminate, haploid gametophyte with a free-living seed plant-like indeterminate, diploid sporophyte. Therefore, the fern life history can be considered a transition between the two other major groups of land plants. The fern reproductive life history offers a unique combined strategy: the indeterminate sporophyte grows at the exact same position as the gametophyte. Dispersal only takes place in the sporophytic generation after the formation of a complex body plan. The absence of dispersal in the fern lifecycle before establishing a complex body plan makes it an interesting model to study evolutionary adaptations to support a nurturing strategy and to explore conserved and divergent properties in land plant embryogenesis. Furthermore, the fern embryo represents the first manifestation of a situation where development of an indeterminate embryo can be guided by maternal tissues. Here, we study embryogenesis in the fern *Ceratopteris richardii* and identify a developmental mechanism that allows for the inheritance of body axis positioning of the few sporophytes that are generated by a gametophyte, presumably favoring progeny establishment in situ.

### High-resolution 4D description of embryogenesis

In the fern *Ceratopteris*, embryogenesis happens in specialized egg-chambers (archegonia) on the haploid gametophyte (Fig. 1A), which produces a heart-shaped hermaphrodite prothallus (*14*) with a meristematic notch where the female sexual organ, the archegonium, develops (*15*). The archegonium (Fig. 1B,C) contains a basket-shaped egg cell (Fig. 1D), supported by subtending basal cells and overlying neck canal cells that form the entry point of the motile sperm (reviewed in (*16*)). Prior to egg maturation, ventral canal cells and neck canal cells block sperm entry. The archegonium is derived from a single cell close to the meristematic notch and develops in a predictable fashion, with clear cell morphologies surrounding the egg cell on the ventral, cross-sectional and dorsal side (Fig. S1). The egg cell induces neighboring cells to divide asymmetrically to form a ring of cells around it, similar to jacket cells in bryophytes. After fertilization the ring of cells is referred to as the calyptra (*14*).

**Fig. 1.**
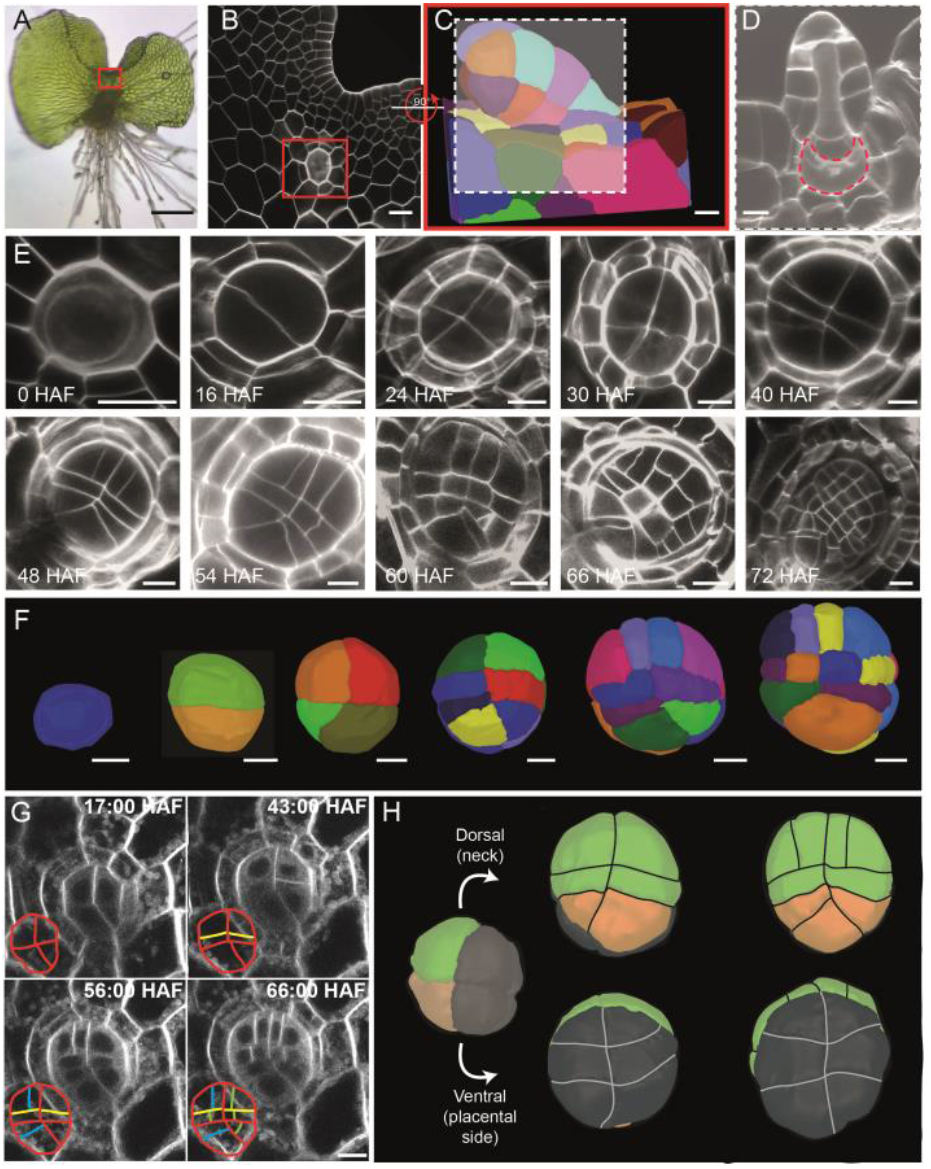
4D characterization of embryogenesis in the fern *Ceratopteris richardii*. (**A**) Sexually mature hermaphrodite gametophyte of *Ceratopteris*. (**B**) Confocal cross-section of the meristematic notch (corresponding to red boxed area in (**A**)). The archegonium is indicated by a red square. (**C**) 3D reconstruction of an archegonium with the dashed box indicating the section visible in (**D**), which represents a longitudinal cross section of an archegonium with the egg cell outlined (dashed). (**E**) Confocal cross-sections representing successive embryonic stages from egg cell to late globular. Time (Hours After Fertilization – HAF) is indicated. (**F**) 3D segmented embryos. (**G**) Stills of live imaging of a single embryo at indicated time points. The time counter indicates hours and minutes after the start of obserbation. (**H**) Dorsal-ventral development of the embryo with the future foot region colored in dark grey. Scale bars are in (A) 200 μm, (B) 25 μm, (C) 10 μm, (D) 10 μm, (E) 25 μm, (F) 15 μm, (G) 20 μm.

Previous descriptions of fern and *Ceratopteris* embryogenesis suggested that there are defined stages of embryogenesis (*17–21*), but no 3D information or lineage inference is yet available. We therefore systematically imaged embryos at different time points following submergence-induced fertilization (Fig. 1E), and used a 3D segmentation strategy to generate a description of embryogenesis (Fig. 1F). We recorded a highly regular succession of cell division events: The first two rounds of division are in the plane of the prothallus (perpendicular to the dorsal-ventral axis), generating a 4-cell embryo. Next, cells divide along the dorso-ventral axis (Fig. 1H), creating an embryo proper (i.e. future embryonic leaf and root) on the dorsal side (archegonial neck) and a foot (i.e. placenta-like structure) on the ventral side (archegonium basal cells) (Fig. S1). A key feature of the embryo is the development of a triangular cell at one end of the embryo, closely resembling the root apical cell of the mature root (*17*), which can be traced back to the basal tier of the four-cell stage (Fig. 1E,F). We confirmed these division patterns in a second *Ceratopteris* ecotype and varied the growth media to rule out asexual embryogenesis (Fig. S2), reported previously in *Ceratopteris* (*22*). Normally, *Ceratopteris* produces a single embryo per gametophyte, by applying sucrose in the growth medium, multiple embryos develop per gametophyte, circumventing this limiting factor. We confirmed our inferred succession of divisions and lineages through live-imaging of embryo development (Fig. 1G, supplementary movies S1-S3). In mammalian embryos, the initial rounds of cleavage divisions partition the cell volume into ever-smaller cells (*23, 24*), and this seems to be shared with *Arabidopsis* (*25*). In *Ceratopteris*, however total embryo size grows continuously throughout development (Fig. S3).

In summary, *Ceratopteris* embryogenesis displays an invariant division pattern similar to seed plants (*25, 26*), which suggests early pattern formation.

### Early patterning and cell differentiation in the Ceratopteris embryo

The regular cell division patterns allow distinct stages to be identified and grouped according to developmental landmarks (Fig. S4; Fig. 2A). We systematically explored embryonic stages to identify such landmarks. Interestingly, in contrast to seed plant embryos (*27*), *Ceratopteris* embryos undergo direct differentiation of cell types. These include the root cap, elongated vascular cells and stomata (Fig. 2B-F). Using these landmarks, we ordered embryos in a developmental timeline and next used RNAseq-based transcriptome profiling to describe the genetic landscape of embryonic development. Embryos were manually collected from 5 different stages, between 2 and 6 days after fertilization (Fig. S5), and RNA-seq was performed. Technical limitations prohibited manual collection for earlier stages (<50 μm). Extensive genetic analysis of *Arabidopsis* embryogenesis has identified many developmental regulators and cell lineage markers (*26, 28*). By comparative genomics, we identified sets of orthologues in *Ceratopteris* and plotted their expression levels across stages (Fig. 2G). The expression levels of these orthologues seem to correlate with the morphological changes during embryogenesis, with vascular and stomata-related genes being enriched in later stages whereas, based on gene expression, ground tissue appears to be established early. Leaf and shoot-related genes appear to be expressed prior to visible organogenesis and root-related genes are also expressed from early stages onward.

**Fig. 2.**
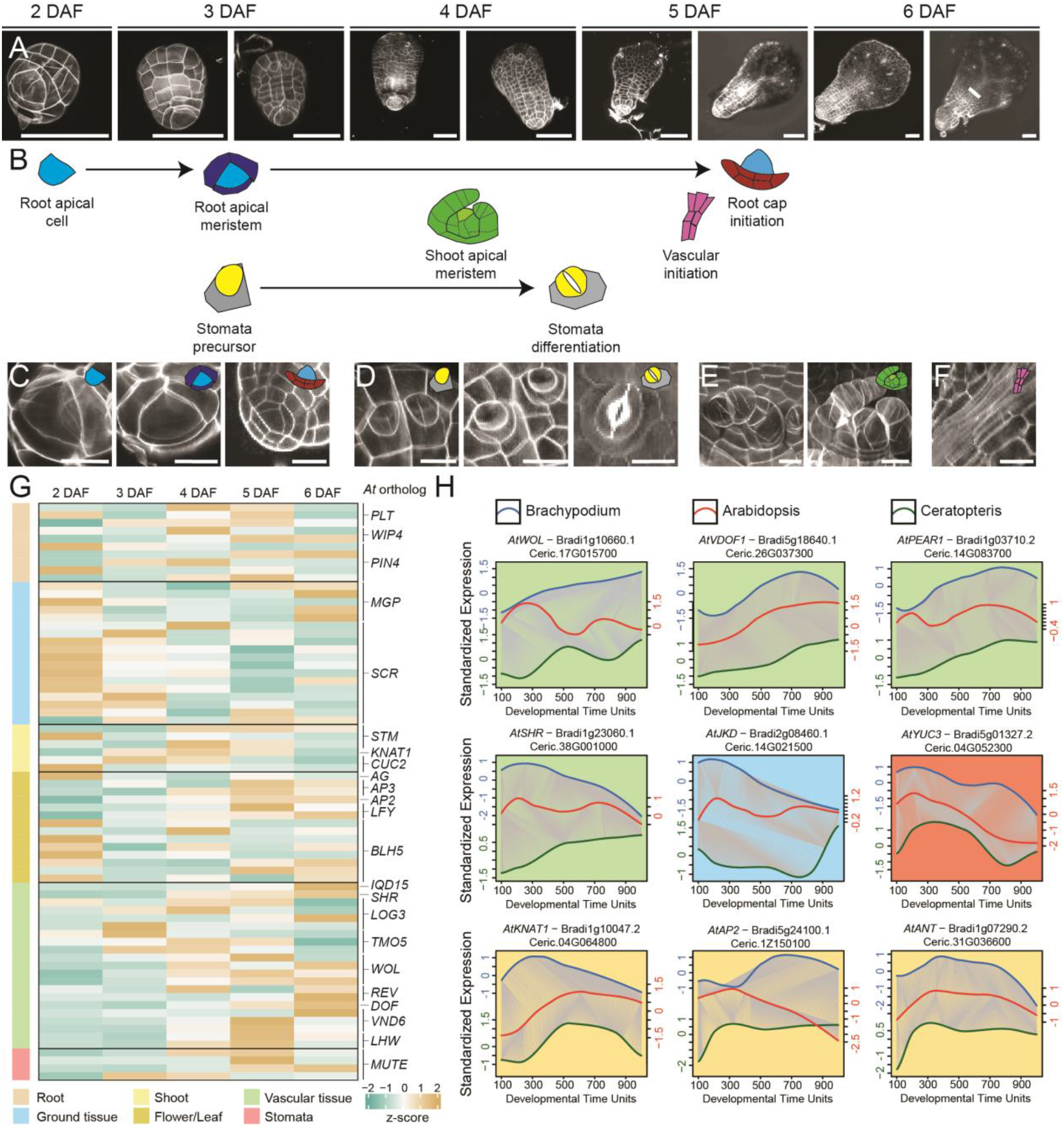
Transcriptional dynamics during *Ceratopteris* embryogenesis suggest early pattern formation. **(A)** Dorsal views of manually isolated embryos used for RNA-seq with (**B**) inferred morphogenetic event and close-ups of (**C**) root patterning, (**D**) stomatal patterning and differentiation, (**E**) shoot apical meristem formation and (**F**) vascular tissue formation. (**G**) Heatmaps of RNAseq-based expression levels of *Ceratopteris* orthologues of *Arabidopsis* developmental regulators and/or patterning markers during embryo development. (**H**) Dynamic time warp analysis of selected *Arabidopsis* embryo patterning genes and closest *Brachypodium* and *Ceratopteris* orthologues. Green shading represents vascular tissue regulators, blue shading ground tissue, red shading auxin biosynthesis and yellow shoot meristem/leaf related genes. Scale bars in (**A**) are 100 μm and in (**C**-**F**) 25 μm.

To further our understanding of transcriptional dynamics, and compare this to seed plants, dynamic time warping (DTW) was performed (*26*) with *Arabidopsis, Brachypodium* and *Ceratopteris* genes. From pre-globular stages onwards, expression was compared across developmental time points for tissue-specific genes known from *Arabidopsis* (Fig. 2H, Fig S6). DTW was not possible for all genes of interest due to the lack of a direct orthologue or a gene not being significantly differentially expressed across either of the three species. In general, temporal expression profiles match those from *Arabidopsis, Brachypodium* or both. Thus, the expression of patterning markers supports the notion that patterning occurs very early during *Ceratopteris* embryogenesis, and that, based on a small set of orthologs, progression of embryogenesis follows similar profiles across vascular plants.

### Maternal control of asymmetric zygote division

Patterning often involves asymmetric cell divisions (ACD) that generate intrinsically different daughters, as well as oriented cell divisions to position daughter cells differently relative to the cellular context (*29*). Asymmetric divisions in the *Ceratopteris* embryo have been described during root apical meristem establishment in the later stages of embryogenesis (*17, 20*), but it is unclear to what extent ACDs may mediate early embryo patterning. We quantified the volumetric asymmetry of the first zygotic cell divisions as a proxy for ACD. Zygote division is asymmetric, generating daughter cells with the larger daughter cell being 40% larger in volume (Fig. 3A,B). This asymmetry seems pre-mitotic, as the volume difference persists over a 6-hour period following zygote division (Fig. 3C). Subsequently, the smaller daughter cell again divides asymmetrically to generate the triangular root precursor cell, while the larger daughter divides symmetrically (Fig. 3D,E). The difference in fate between zygotic daughters is also reflected in a distinct difference in vesicle density between apical and basal tiers from the four-cell stage onwards (Fig. 3F). Thus, the first two rounds of division in the embryo are highly asymmetric both in geometry and in fate, and they generate the root pole in an extremely early patterning event.

**Fig. 3.**
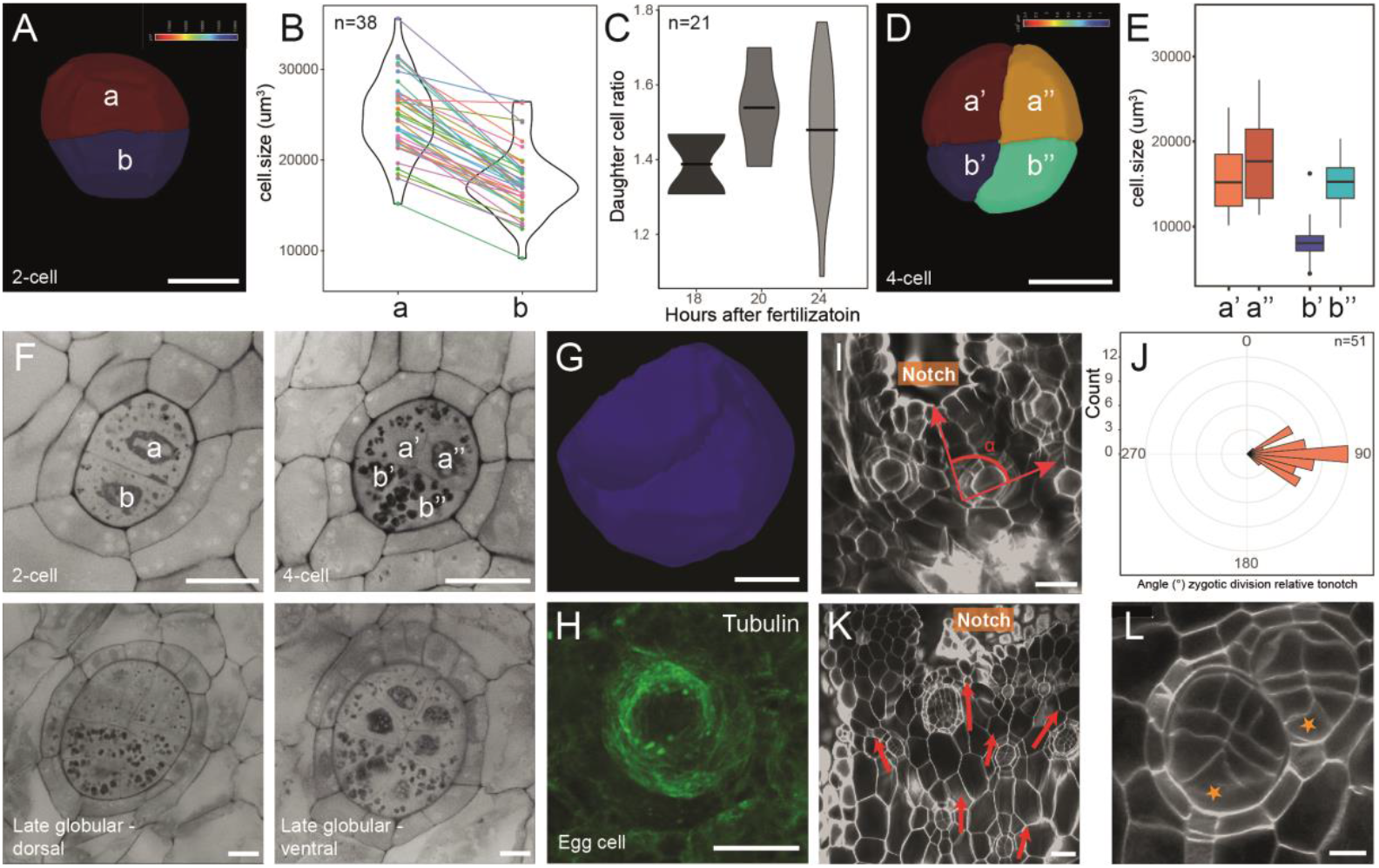
Early axis establishment in the *Ceratopteris* embryo is correlated with maternal tissue morphology. **(A)** 3D volume segmentation of a two celled embryo with ‘a’ depicting the apical sister cell while ‘b’ depicts the more basal sister cell. (**B**) Volumes of the two different daughter cells in two-celled embryos with colored lines depicting the two daughters of a single embryo and (**C**) quantification of the volume ratio between sister cells at different developmental time points. (**D**) 3D volume segmentation with quotation marks depicting the different daughter cells and (**E**) quantification of the volume of the different daughter cells. (**F**) Cytosolic counterstain of cleared embryos from different developmental stages to show cellular differentiation. (**G**) 3D egg cell morphology and (**H**) microtubule immunostaining of a mature egg cell. (**I**) Orientation of the zygotic division with respect to the maternal meristematic notch with red arrows depicting the angle measured to quantify the angle between the zygotic division and the direction of the maternal notch meristem. (**J**) Quantification of the angle of the zygotic division relative to the meristematic notch with 90° meaning that the zygotic division is perpendicular to the axis to the notch meristem. (**K**) Example of multiple embryos on a single gametophyte, all showing the same axis of development with the asymmetric divided daughter cell at the basal site of the embryo, red arrows show the orientation of the embryo. (**L**) A twin embryo/archegonium with a shared direction of development. Scale bars are in (A) 20 μm, (C) 30 μm (F) 25 μm, (G) 10 μm, (H) 15 μm, (I) 30 μm, (K) 50 μm and (L) 20 μm.

The zygote divides asymmetrically, and the question is what guides this asymmetry. The egg cell itself is dorsoventrally polarized due to its basket-like shape (Fig. 3G; (*18–20*)) but this axis is unrelated to the future ACD axis (Fig. 1B,C and D). Additionally, egg cell morphology changes dramatically after fertilization, towards a sphere (*18*). Microtubule arrangement in the egg cell aligns with the distinct shape of the egg cell, but shows no clear correlation with the future zygote ACD (Fig. 3H). We therefore infer that zygote ACD is not guided by intrinsic egg cell polarity. However, we found that zygote division is highly oriented within the maternal tissue (Fig. 3I, J), as has previously been suggested (*18, 20*). We find that this positions the larger (apical) daughter cell closer to the notch meristem, and leads to a fixed embryonic axis with the root pole pointing away from the meristematic notch, and towards the rhizoids (Fig. 3K; Fig. 1A). This orientation seems independent of archegonium structure as embryo axis orientation is maintained in occasional twin embryos (Fig. 3L). These data suggests that zygote ACD is correlated with maternal tissue morphology, and that the alignment of the sporophyte shoot-root axis is aligned with the maternal meristem-rhizoid axis.

### Symplastic isolation of the early Ceratopteris embryo

The ACD-directing influence of the gametophyte on the *Ceratopteris* zygote can be of any nature. Given that soluble compounds such as plant hormones (e.g. auxin) can influence flowering plant embryo development (*30–33*), we explored whether chemical communication may underlie maternal control of embryo polarization. We first asked whether there are symplastic connections between the gametophyte and the embryo. Transmission electron microscopy (TEM) showed the presence of a clear cavity between the embryo and maternal gametophyte tissue (Fig. 4A), and cell wall thickenings (Fig. S7) where the primary cell wall seems to consist mainly of cellulose (Fig. S8). While the two generations were clearly isolated in the plane of axis polarization, in contrast we confrimed plasmodesmata, and therefore symplastic connections, between embryonic cells and between gametophyte cells (Fig. 4B,C). Furthermore, we detected the presence of an envelope-like structure (Fig. 4D,E; Fig. S9), likely homologous to the recently identified *Arabidopsis* embryo envelope (*34*). Taken together, this suggests that there is no connection between the embryo and the maternal tissue in the future axis of polarization and that the embryo is surrounded by a physical barrier.

**Fig. 4.**
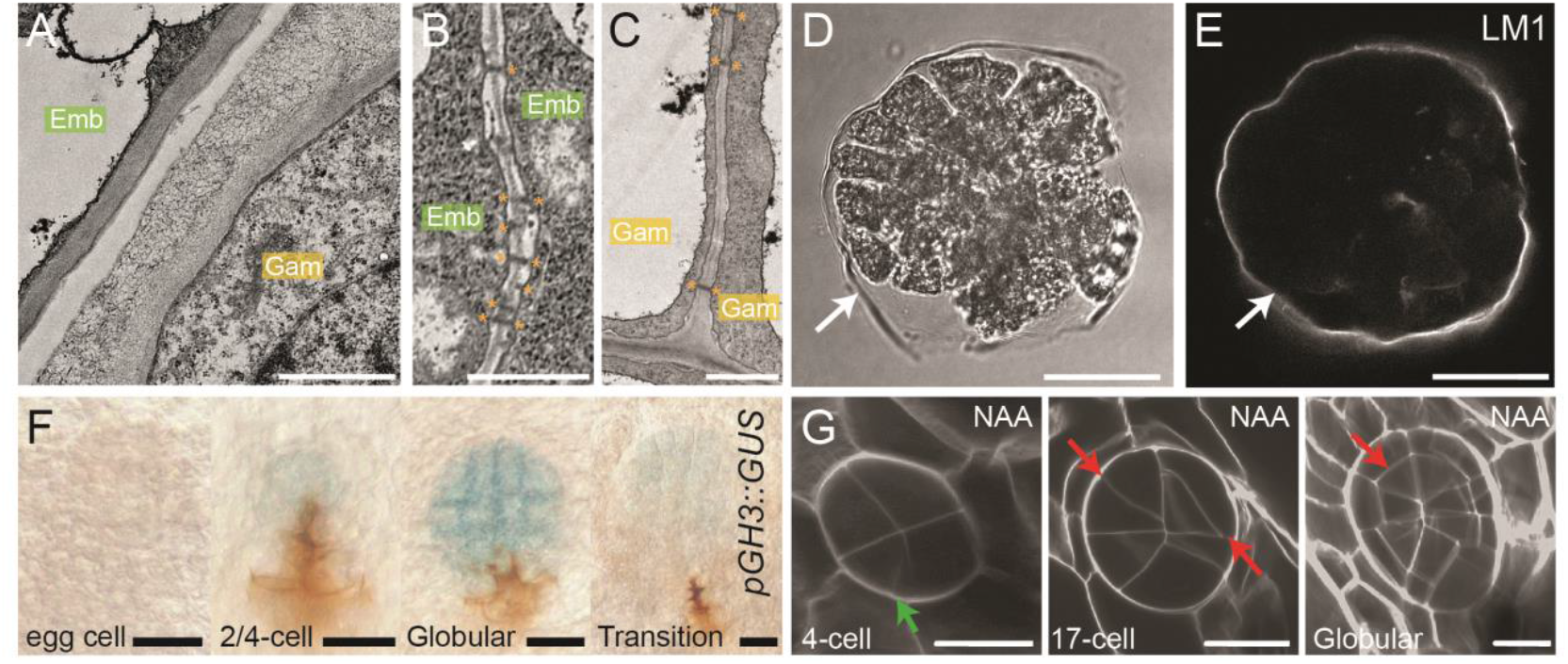
Limited chemical signaling and the lack of auxin-effects on zygote polarization. **(A)** TEM image showing the boundary between the embryo (Emb) and maternal gametophytic tissue around it (Gam). (**B**) Plasmodesmatal connections between embryonic cells and (**C**) between gametophytic maternal cells. (**D**) Bright field image of a manually isolated embryo with a digested cell wall and the same embryo in (**E**) stained with the LM1 antibody that detects Extensin epitopes. White arrows in (D) and (E) point at the envelope-like structure. (**F**) GUS staining of different embryonic stages of the auxin reporter pGH3::GUS, (**G**) Exogenous application of synthetic auxin and its effect on different embryonic stages with green arrows depicting normal divisions while red arrows depict abnormal divisions. Scale bars are in (A) 1 μm, (B) 1 μm, (C) 1 μm, (D) 40 μm, (E) 40 μm,, (F) 25 μm and (G) 30 μm.

To further investigate the role of chemical communication in embryo polarization, we turned to auxin, based on its undisputed importance during *Arabidopsis* embryo patterning (*30–33*). We generated a transgenic line carrying the auxin-responsive pGH3::GUS construct ((*35*); Fig. S10;S11). We detected GUS activity from the 2/4-cell stages onward, yet we observed no clear local activity until later stages (Fig. 4F; Fig S12). This suggests that the auxin response is active in embryos, but that there is no clear response output in the egg cell or zygote. We next aimed to manipulate auxin activity by treating plants with exogenous auxin before and at the moment of fertilization. This did interfere with later embryo divisions, but did not affect the initial zygote ACD or the second round of divisions (Fig. 4G, Fig S13). Hence, while auxin response is active in the embryo, external manipulation of auxin accumulation does not change early embryo polarization. Together with the symplastic embryo isolation, this suggests that maternal control of zygote ACD may not be mediated by chemical signals.

### Tissue mechanics as a patterning cue for zygote ACD

Besides chemical cues, development in multicellular eukaryotes uses cellular and tissue mechanics as instructive signals (*36, 37*). Mechanical stress guides oriented divisions in the *Arabidopsis* shoot and sepal, can affect expression of homeotic genes (*38, 39*), helps orient stomatal divisions in the leaf (*40, 41*), and orient cambium cells in the vasculature (*42, 43*). Given the correlation of zygote ACD with the fundamental gametophyte body (notch-rhizoid) axis, we asked whether tissue mechanics may couple polarities across generations. We therefore developed a finite element model (FEM) based on realistic gametophyte shapes (Fig. 5A). From confocal stacks, the cell wall network was extracted (Fig. S14A-E) and used as a basis for modeling. The tissue was modeled as a quasi-2D structure, as most of the prothallus is only a single-cell layer thick, with the archegonium as an exception. Modeling showed that an anisotropic stress pattern (see material and methods) can emerge with a maximum tension that correlates with the direction of the zygotic division (Fig. 5B), if a gradient in cell wall stiffness from the meristematic notch (i.e., dividing cells) to the differentiated basal part is assumed (Fig. S14N, O; see Supplementary materials). The predicted stress pattern was insensitive to the shape variability among 8 individual plants (Fig. S14F-M). The assumption of differential cell wall stiffness between dividing (notch) and growing (thallus) cells is in line with atomic force microscopy data collected in the *Marchantia* thallus (*44*).

**Fig. 5.**
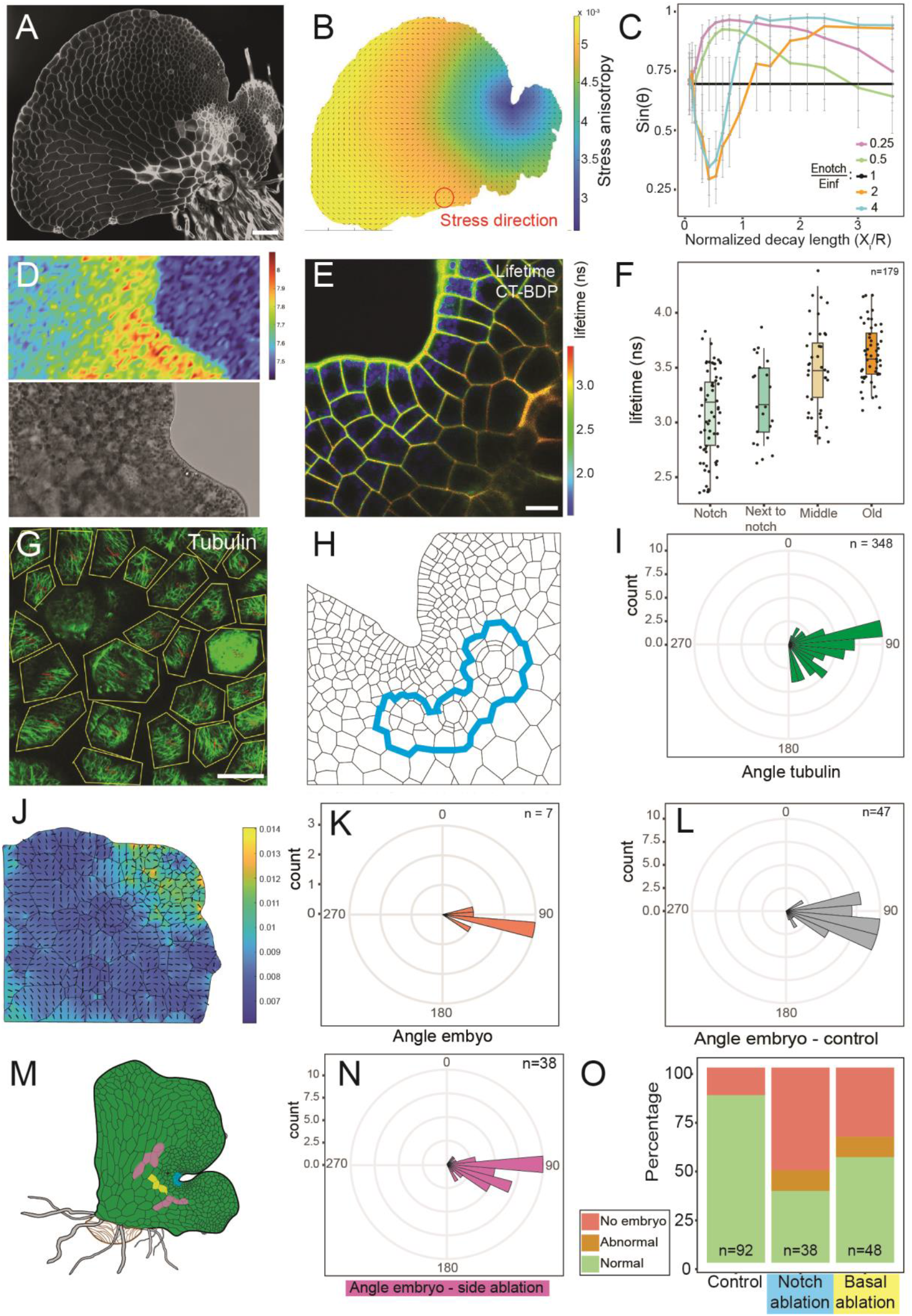
Mechanical stress fields correlate with zygotic division and are dependent on meristem organization. **(A)** Microscopic image of the cell wall network of sexually mature gametophyte and the corresponding **(B)** modelled stress anisotropy and direction of maximum tension. (**C**) Modelled angles of egg cell stress anisotropy under different assumptions regarding the stiffness gradient (E_notch_/E_∞_) and the decaylength of this gradient ξ/ relative to the distance R from the notch to the archegonium, error bars represent standard error. Angle is depicted as sin(θ) and is the same angle as shown in Figure 3I/J. If sin(θ) equals 1 it represents the angles (±90°) measured in figure 3 (**D**) Brillouin microscopy of the meristematic notch with the top panel showing the Brillouin shift and the bottom panel the accompanying brightfield picture of the same plant. (**E, F**) FLIM picture depicting the lifetimes of the carbotag-bodipy probe (E) and quantification of lifetimes from multiple individual plants with cell walls grouped according to their distance from the notch (F). (**G**) Example of FibrilTool output to quantify orientation of microtubules. (**H, I**) Schematic representation of the cells for which the orientation of microtubules was quantified related to the notch (H) and the quantification of microtubule orientation in those cells (I). (**J**) NAA grown gametophyte and its modelled stress anisotrophy. (**K**) Orientation of embryos relative to the notch meristem of NAA grown plants as shown in (J). (**L**) orientation of control embryo’s relative to the notch meristem. (**M, N, O**) Surgical ablation of different groups of cells of the gametophyte prior to fertilization with color coding depicting the ablation sites. (M), orientation of two-day old embryos which developed after gametophytes where ablated at their lateral wings (N) and percentage of gametophytes developing embryos when ablated at the notch or basal site below the archegonia prior to fertilization (O). Scale bars are in (A) 75 μm, (D) 30 μm, (E) 15 μm and (G) 30 μm.

We further explored the requirements of the stiffness gradient for zygote division angles. The modelling suggested that the gradient should have a >2-fold stiffness difference between walls at the notch meristem and the basal walls, and a decay length that is similar to the distance from the notch meristem to the archegonium (Fig. 5C). We next used two approaches to map wall stiffness in prothallus. Firstly, Brillouin microscopy reported a robust gradient in frequency shift (Fig. 5D, Fig. S15). The frequency shift detected with Brillouin microscopy allows calculation of the longitudinal modulus, which can be considered a proxy for stiffness (*45*). The measurements suggest that stiffness levels are highest in the notch area and are getting lower while cells are differentiating. Secondly, a fluorescent cell wall microviscosity probe (*46, 47*) revealed differences in internal cell wall viscosity – an independent proxy for stiffness – at the same length scale (Fig. 5E,F). Neither of these methods allows for a calibrated inference of actual stiffness, but both *in vivo* methods show a gradient in mechanical properties with decay length in the same order as predicted by the FEM modeling to guide zygote ACD. We therefore conclude that tissue-derived stress fields correlate with zygotic division orientation.

Mechanical stresses influence the orientation of the microtubule (MT) cytoskeleton, which aligns with maximal tensile stress (*48, 49*). Given the critical role of MT in guiding cell division orientation, they are a potential coupling factor between mechanics and ACD orientation. Immunostaining of MT in the *Ceratopteris* gametophyte showed a strong correlation of MT orientation with the modeled direction of tensile stress in the cell files surrounding the archegonia (Fig. 5 G, H and I). In most cell files, MT orientation corresponded to the modeling, except for the outer cell layer and central region of the notch meristem (Fig. S16). We probed the relevance of the MT cytoskeleton for zygote ACD and embryo development by treating gametophytes with the MT depolymerizing drug oryzalin prior to fertilization (Fig. S17). Whereas high oryzalin concentrations led to the absence of zygotic division and swelling of the zygote, low concentrations induced embryo malformation (Fig. S17).

The mechanism that emerges is that tissue stresses that are generated by juxtaposition of growing and dividing cells with different stiffness instruct MT orientation and cell division. This mechanism should act on all dividing cells, and would not be selective for the zygote. We asked whether this is the case by mapping patterns of cell division through EdU staining. Most cell divisions take place on the outer cell layers, except the specialized division forming the archegonium (Fig. S18A). Thus, zygotes represent the few cells away from the notch that actually divide. The zygote is subtended by a placental cell, which should behave similarly to the zygote, given its exact same position in the stress field. Indeed, placental cell division correlated strongly with zygote division (Fig. S18B,C), suggesting that the same principles guide division orientation and that triggering a division in this region leads to oriented division regardless of cell identity.

Lastly, we tried to experimentally manipulate the system to derive causalities. The fragility of single-cell layer-thick prothallus and the absence of genetic tools make it challenging to perturb development and mechanics. We turned to two perturbation strategies. First, we asked what the requirements are for thallus tissue geometry. Growth on synthetic auxin leads to a deformed prothallus shape with a rounded and elongated shape instead of a cordate shape (*50*). Modeling these shapes suggests an unperturbed stress field (Fig. 5J, Fig. S19). In these prothalli, we found that embryo orientation was indeed normal (Fig. 5K,L), suggesting that the mechanical stress field requires only a notch-rhizoid axis. Next, we used needle-induced surgical ablations of small regions surrounding the archegonium to disrupt tissue integrity prior to fertilization. Consistent with the result of NAA treatment, ablation of the gametophytic “wings” does not affect embryo orientation (Fig. 5M,N). However, surgical ablation of the notch meristem or cells between the archegonium and the rhizoids caused to some extent embryonic arrest (i.e. lack of embryos) and malformed embryos (Fig. 5O, Fig. S20). These manipulation experiments support the notion that the integrity of a notch-rhizoid axis is a minimal yet critical requirement for zygote ACD.

## Discussion

By studying embryogenesis in the fern *Ceratopteris*, we discovered a nurturing mechanism that we interpret to be an adaptation to the reproductive strategy in which normally a single, indeterminate embryo is generated by maternal gametophyte tissue. Key to this mechanism are two features: (1) a highly regular cell division pattern that generates the future root pole within two rounds of division and (2) a coupling between maternal tissue geometry and the orientation of the first, asymmetric zygote division. Regular embryonic cell divisions are found in other plant species (*51, 52*), but it is unclear whether there is a strict requirement for such patterns, given the impressive regenerative potential most plants display (*53*). Whether critical or not, the extremely early deterministic division pattern in *Ceratopteris* embryos does position the root pole and thus determine the shoot-root axis. At present, we do not know what mechanisms drive the regular division patterns, but it is possible that – as in *Arabidopsis* – patterning on a template with relatively few cells imposes a constraint that limits variability. Given that archegonia are the ancestral state of reproductive structures in land plants (*54*), this form of embryogenesis may well reflect an ancestral state of embryogenesis in all seed plants. Perhaps the alignment of the *Arabidopsis* zygote division with maternal tissue polarity (*25*) is a remnant of shared principles among vascular plants. We expect that the further development of genetic tools in *Ceratopteris* and the study of related ferns may help address these questions.

We find that the orientation of the first zygote ACD is strongly biased by the gametophyte notch-rhizoid axis, and as a consequence, the embryonic shoot-root axis aligns with it. We explored the role of chemical communication in guiding the orientation of the zygote ACD, but find that symplastic isolation and the presence of a barrier may limit such influences. Indeed, we could not find evidence for an instructive role of auxin, whereas such a transgenerational patterning influence has been reported in flowering plants (*30, 32*). Clearly, we have not exhaustively explored further chemical signals – including signaling peptides (*55*), and we cannot exclude that the placental connection allows the generation of signal gradients in the overlying embryo tissue.

Regardless of the unknown role of chemical signals, we find that zygote ACD orientation is guided by mechanical stress patterns that evolve as a consequence of the gametophyte structure. Both modeling and experimental data show that potentially instructive stress patterns can form with a minimal requirement of the juxtaposition of meristematic notch and growing prothallus cells. These stress patterns are therefore a robust mechanism that – when coupled to the mechanical sensitivity of MT and the local positioning of the egg cell, can explain the orientation of the first division. Similar mechanical interactions have been described in the *Arabidopsis* seed (*56*). However, this mechanism cannot explain the asymmetry of the first division. There is thus a clear need to identify markers for cell polarity (such as SOSEKI; (*57, 58*)) to address cellular and molecular mechanisms in the future.

We stress a limitation of this work in that it is very challenging to firmly establish causality. All results are consistent with a key role for tissue geometry, but the development of tools to selectively manipulate the stress pattern will be needed to demonstrate sufficiency in guiding ACD orientation. Nonetheless, it is intuitive that positioning the root towards the functionally analogous rhizoid pole would enhance successful sporophyte establishment, and tools to manipulate the ACD orientation will help derive the ecological significance of axis inheritance.

## Acknowledgments

We thank the Wageningen Electron Microscopy Center (WEMC) for generating the TEM images and the Microspectroscopy Research Facility (MSC) for giving acces to their facilities. We did not make use of any editing or AI-assisted technologies. We thank R. Smith for helpful assistance with the 3D segmentation, M. Nodine (Wageningen University) for technical advice for RNA-library preparations, D. dos Santos Pereira (Wageningen University) for bioinformatic help, M. Besten (Wageningen University) for help with the Carbotag probes, J. J. Ramalho (Utrecht University) for support with starting up the project, Jelmer Vroom (Wageningen University) for the TEM imaging, J. Rudolph for his assistance in the project, Hisa Yoshida, Junko Kato and Yusuke Kimata for providing materials for live imaging and Ester Rabbinowitsch (Oxford University) for help with generating transgenic lines.

## Funding

This work was supported by a grant from the Graduate School Experimental Plant Sciences to S.W., by the Netherlands Organization for Scientific Research (NWO; OCENW.KLEIN.516) to D.W. and J.S., a Gravitation grant (GreenTE) of the Dutch Ministry of Education to D.W., J.v.d.G. and J.S..

The European Research Council, ERC AdG (EDIP) to JAL.

The National Natural Science Foundation of China (Grant No. 32471892) to Z. H.

European Research Council, ERC HYDROSENSING project 101118769; Faculty of Natural Sciences, Norwegian University of Science and Technology (NTNU), Natural sciences (NV) faculty, Career Development grant to L.A.B.; Brillouin microscope building grant from NV faculty at NTNU. Norwegian Research Council WALLINTEGRITY project 315325 to T.H. The Japan Advanced Plant Science Network, the Japan Society for the Promotion of Science [a Grant-in-Aid for Research Activity Start-up (JP21K20649 to H.S.), a Grant-in-Aid for Early-Career Scientists (JP24K18135 to H.S.), a Grant-in-Aid for Scientific Research (B) (JP23H02494 to M.U.), and International Leading Research (JP22K21352) to M.U.], the Japan Science and Technology Agency [CREST (JPMJCR2121 to M.U. and YORC to H.S.)], the Yamada Science Foundation (to H.S.), the Suntory Rising Stars Encouragement Program in Life Sciences (SunRiSE to M.U.), and the Toray Science Foundation (20-6102 to M.U.).

## Author contributions

Conceptualization: SW, DW

Methodology: SW, ZH, HS, LAB, CB, AP

Investigation: SW, ZH, HS, LAB, CB, AP

Modelling: JvG

Visualization: SW, JvG, ZH, DW

Funding acquisition: DW, JS, JvG, JAL

Supervision: DW, JS, MU, TH, JAL

Writing – original draft: SW

Writing – review & editing: DW, SW, with edits from all coauthors

## Data and materials availability

The RNAseq raw reads have been deposited in the NCBI Short Read Archive (SRA) under the BioProjectID PRJNA1293105. All transgenic lines are available upon request. All microscopy data is available upon request. All quantifications are available in the supplementary materials.

## Supplementary Materials

Materials and Methods

Figs. S1 to S21

References(*59*-79)

Movies S1 to S2

Data S1

