## Supplemental File for "Transgenerational polarity axis inheritance during *Ceratopteris* embryogenesis"

**The PDF file includes:**

Materials and Methods
Figs. S1 to S21
Table S1
References 59-79

**Other Supplementary Materials for this manuscript include the following:**

Movies S1 to S2
Data S1

### Materials and Methods

#### Plant material growth conditions

Spores of *Ceratopteris richardii* (strains Hn-n and 176D (*59*)) were grown according to Plackett, Huang, Sanders and Langdale (*60*) in a Hettich MPC600 incubator set at 28°C, with 16 h of 100 µmol $m^{-2}$ $s^{-1}$ white light. Plants were grown on ½-strength MS medium with 1% sucrose to induce the formation of multiple embryos per gametophyte. Gametophytes were grown until sexual maturity (± 10 days) when the notch meristem was established and archegonia were visible. For fertilization, plants were flooded with a surplus of demineralized water (>10 mL per plate).

#### Microscopy - Cell wall staining and clearing

For embryos attached to gametophytes, entire prothalli were collected and cleared with ClearSeeAlpha for at least 4 days while rotating (*61*). Embryos were collected at different timepoints after fertilization (induced by flooding – taken as timepoint zero). For archegonia imaging, immature gametophytes were collected at different timepoints of development. After clearing, plants were stained in ClearSeeAlpha with 0.1 % Renaissance SR2200 (*62*) for 1 hour to over-night. Imaging was performed on a Leica SP5 confocal microscope with standard settings for Renaissance SR2200 (405nm excitation, 425-460nm detection (*62*)). For 3D segmentation, image settings were changed to 16-bit and z-stacks were taken with step sizes that approached cubic voxel sizes. Z-steps were kept between 0.1 and 0.25 µm to ensure proper 3D segmentation.

#### Microscopy - 3D segmentation

3D segmentation of embryonic z-stacks was done using MorphographX (https://morphographx.org/\). 3D segmentations were always visually checked and manually curated. In general, Z-stacks were loaded into the software. Voxel sizes were adjusted to the used image settings. Stacks were blurred using the Gaussian blur function (set at 0.3-0.5) and contrast was subsequently enhanced (Brighten/Darken set at 2-16). 3D segmentation was done using the autoseeded watershed algorithm (set at 500-2000). Segmentation was repeated with different parameters until the results reflected the loaded stacks. Lastly, over-segmented cells were manually fused together. A 3D mesh and its cell shapes were subsequently extracted using a modified marching cube algorithm (1-4 µm). 3D meshes were subsequently used to extract cell volumes and compute division analysis, as been described previously (*25*).

#### Microscopy - Embryo cytosolic counterstaining and archegonium cell wall staining

Cleared embryos were stained with Nile Red (oxazone) and not washed prior to imaging. Normally, Nile Red stains lipids, but in cleared samples, it also stains the cytosol. For archegeonia, cleared gametophytes were stained with different cell wall dyes as described by **Ursache, Andersen, Marhavý and Geldner (*63*)**. Imaging was done with a Leica SP5 system

#### Microscopy - Tubulin staining and analysis

Microtubule visualization was done by immunostaining of tubulin in sexually mature gametophytes. Fixing of the tissue was done according to Apostolakos and Galatis (*64*) without mounting the tissue on a cover slip. Enzymatic digestion was performed with 1.5% Cellulase Onozuka R-10 (Duchefa), 1% Driselase (Sigma) and 1% Macerozyme R-10 (Duchefa). Immunostaining was performed by diluting the anti-α-tubulin (DM1A)-Alexa Fluor™ 488 antibody (eBioscience™, Invitrogen) 1:100 in PBS with 1% BSA. Staining was done overnight

at room temperature. Before imaging, samples were washed once in PBS with 1% BSA. Imaging was done on Leica SP5, Leica SP8 and Leica SP8 DIVE multiphoton/2-photon systems. 2-photon microscopy was used to image microtubule orientation within the egg cell.
Microtubule orientations were quantified using the FibrilTool with default parameters (***65***). Subsequent orientations of microtubules were manually quantified in ImageJ to align to the notch meristem.

**Microscopy - Live imaging**
To visualize early embryos, gametophytes were stained as described previously (*66*). In brief, gametophytes were kept in the liquid *in vitro* ovule cultivation medium (*67-69*) containing 10 µM of FM4-64 (BioTracker 640 Red C2; Sigma #SCT127) for several hours. For microscopy, the pre-stained gametophytes were held in place on a 35 mm diameter glass-bottom dish (Matsunami Glass Ind., Ltd., Osaka, Japan) filled with the aforementioned liquid medium containing FM4-64, using a 0.2 mm-thick silicone sheet. Live imaging was performed as described previously (*66*) using a Nikon A1 inverted laser-scanning microscope equipped with Ti:sapphire femtosecond pulse laser (Mai Tai DeepSee; Spectra-Physics, California, U.S.A.). Fluorescence signals were detected by the external non-descanned GaAsP photomultiplier tube detectors. A dichroic mirror (DM560) and a standard mirror were employed to direct the excitation and emission light paths. Embryo images were acquired at 61 z-stacks with 1.5 µm-intervals every hour using a 40× water-immersion objective lens (CFI Apo LWD WI; Nikon, Tokyo, Japan) with Immersol W 2010 (Zeiss, Oberkochen, Germany) as the immersion medium. All visualization procedures of the acquired data were performed using Fiji (https://fiji.sc/), including manual adjustment of brightness and contrast, extraction of single XY planes from each time point, and conversion into a movie file.

**Microscopy - Transmission electron microscopy (TEM)**
Gametophytes which were fertilized two or three days prior were fixed in 2.5% glutaraldehyde and 2% paraformaldehyde in 0.1M phosphate/citrate buffer (pH 7.2) overnight at 4°C. Samples were washed 6 times for 10 minutes with 0.1M PBS. After washing, samples were fixed in 1% Osmium tetroxide in 0.1M PBS for 1h at room temperature. Samples were washed again for three times 10 minutes in MilliQ water. Samples were dehydrated by using an ethanol series (10%, 30%, 50%, 70%,80%,90%,96%,100%,100%), with every step for 15 minutes except the last step at 30 minutes. After dehydration, samples were infiltrated with resin (Spurr'r resin) in a stepwise fashion by increasing the resin: ethanol ratio. 1:2 resin:ethanol for 30 min, 1:1 resin:ethanol for 30 min, 2:1 resin:ethanol for 30 min, and 100% resin for 60 min to overnight at room temperature (Spurr's kit Catalog number 14300) After infiltration, samples were transferred to gelatin (LR white) and polymerized within a capsule for 24 hours at 60-65°C. Polymerized samples were pre-sectioned using a Leica EM RAPID microtome and sectioned into 50-nm-thick sections using a Leica ultramicrotome UC7 and attached to the grid. Grids were incubated for 10 minutes in 2% uranyl acetate. Samples were washed five times with water and subsequently incubated for 10 minutes in ready-to-use lead citrate (EMS) in a $CO_2$-free environment. Samples were washed two times in 0.01N $CO_2$-free water, and afterwards washed three times in MilliQ water. Samples were imaged in a TEM (JEOL JEM1400) at 120kV.

**Microscopy - Embryo envelope imaging**
The extra cell wall structure around the embryo was imaged and stained as described in Harnvanichvech, Borassi, Daghma, van der Kooij, Sprakel and Weijers (***34***). Briefly, embryos

were manually dissected and subjected to enzymatic digestion for 30 minutes. Enzymatic mixture consists of 20 mM MES (pH 5.7) containing 0.4 M mannitol, 20 mM KCl, 2% (w/v) Cellulase R10 (Duchefa, C8001), 0.5% (w/v) Macerozyme R10 (Duchefa, M 8002), 0.05% (w/v) Pectinase (Sigma-Aldrich, P2401), 0.05% (w/v) Hemicellulase (Sigma-Aldrich, H2125) and 0.05% (w/v) Driselase (Sigma-Aldrich, D9515). After digestion, embryonic cell walls were stained with 0.1% (w/v) Calcofluor White (18909, Sigma-Aldrich) or Aniline Blue (415049, Sigma-Aldrich). Immunolabeling was done by initially incubating in 20 µl of 3% (w/v) milk protein in 1× PBS (MP/PBS) for 15 min to block non-specific binding sites. Embryos were then incubated with primary monoclonal antibodies (LM1 as18 4210, LM2 as18 4211) diluted 1:10 in MP/PBS for 15minutes and washed afterwards with 1x PBS. For the secondary antibody, embryos were incubated in anti-mouse-IgG linked to fluorescein isothiocyanate (Sigma-Aldrich, F6258) diluted 1:100 in MP/PBS for 15 min

**Microscopy - Brillouin Microscopy**

Brillouin microscopy was performed using a custom-built confocal Brillouin microscopy based on a two-stage imaged phase array (VIPA) configuration, following the design described by **Zhang and Scarcelli (*70*)**. Excitation was provided using a 532nm laser (Cobolt, Hübner Photonics). The laser was guided to the back port of an inverted Leica SP8 confocal microscope. Samples were illuminated using a 40x water-immersion objective NA 1.1 (Leica) and the scattered light was collected using a backscattered geometry. The spectrometer consisted of two-stage VIPA etalons (OP-6721-3371-2, 500-600 nm, 30 GHz free spectral range - Light Machinery) and a Lyot stop **(*71*)**. An ORCA-Quest qCMOS camera (C15550-20UP, Hamamatsu Photonics) was used to record the Brillouin spectra. Images acquisition was performed using a physical pinhole of 1 Airy unit, 100ms acquisition time per frame (with frame integration equal 2), stage step sizes of 1-3mm and laser power 10-15mW. The microscopy room was maintained at 20°C and the laser was mounted on a heated plate set to 37°C (according to manufacturer recommendations). To check for perturbations and phototoxicity, samples were inspected by transmitted-light wide-field illumination before and after the Brillouin spectra was recorded. No visible changes were observed. Water and methanol were used as reference samples to calibrate the frequency axis. During Brillouin spectra recording, the confocal microscope stage movement and image acquisition was controlled using the HCImage software (Hamamatsu Photonics). A custom MATLAB script was used to extract the Brillouin peaks position after fitting the peaks by a Lorentzian function. Peak positions, together with the calibration samples, are used to calculate the frequency shift (frequency difference between the Stokes/AntiStokes peak with respect to the Raileigh peak). The frequency shift is used as a proxy for the elastic behavior of a material as it relates to the longitudinal modulus of by the formula:

$$v_B = \frac{2n}{\lambda}\sqrt{\frac{M'}{\rho}}\sin\frac{\theta}{2}$$

Where nB is the frequency shift, $n$ is the refractive index, $\lambda$ is the excitation laser frequency, $M'$ is the real part of the longitudinal modulus, $\rho$ is the material density and $\theta$ is the scattering collection angle.

Brillouin values are reported as the relative frequency shift of the sample with respect to the frequency shift value of water.

**Microscopy - Cell wall porosity FLIM measurements**

For imaging cell wall porosity with the carbotag-bodipy probe, methods were recently described Besten, Hendriksz, Michels, Charrier, Smakowska-Luzan, Weijers, Borst and Sprakel (*46*). Staining was done overnight in liquid ½ MS at a concentration of 1µM. Imaging was done on a Leica SP8 Dive multiphoton with the associated FLIM module. For every image, an accumulation of 20 was used to ensure enough photons per pixel, while laser intensity was adjusted to prevent multiple photons hitting the detector at the same time. Quantification of lifetimes was done by plotting a 2-component decay curve upon every picture. FLIM lifetimes were extracted by using the second component and manually selecting and classifying cell walls of interest.

**Microscopy - EDU staining of S-phase cells**

Gametophytes of different ages were grown for 4 hours in liquid ½ MS supplemented with 40 µM EdU, followed by fixation in 4% formaldehyde in PBS (PH = 7.2) for 1 hour under vacuum. During these hours, tubes were removed multiple times from vacuum and agitated. After vacuum, samples were washed with PBS and permeabilized with PBS with 0.1% Triton X-100 for 20 minutes. Subsequently, samples were washed twice with PBS with 3% BSA before staining according to the click-IT EDU reaction mixture for 1 hour with the 594 Alexa FLUOR fluorophore (Invitrogen). After staining, samples were washed once with PBS and placed in ClearSeeAlpha. Imaging was done with SR-2200 as a cell wall counterstain.

**Image processing and analysis**

All images were processed in Fiji ((https://fiji.sc/), including manual adjustment of brightness and contrast, extraction of single XY planes and for live imaging, conversion into a movie file. Angles of divisions were manually measured using the angle tool.

**Transcriptome analysis - RNA isolation and sequencing**

For RNA isolation, embryos were manually isolated from the maternal gametophytic tissue at different timepoints after fertilization (induced by flooding). Isolation was done using two microlance injection needles (BD) and a stereoscope (Motic, K500) at max magnification (40x). One needle was used to make an incision next to the archegonium, after which the second needle was used to apply pressure on the other side of the archegonium, often resulting in the bursting of the archegonium and the release of the developing embryo. Embryos were collected using a P10 pipette tip and resuspended in 10% RNAlater (ThermoFisher) on ice. Multiple embryos were collected for each sample (n>30 for the earliest stages and n>15 for the latest stages). After finishing collecting one sample, embryos were frozen at -70°C until further processing. Attention was paid to freeze samples within 1h after collecting the first embryo to prevent RNA degradation. Earlier stages of embryogenesis proved to be impossible to collect manually due to their small size (<50µm).

For RNA isolation. embryos were dissociated in Trizol (Invitrogen). RNA was subsequently extracted using chloroform. RNA was pelleted in isopropanol with the co-precipitant glycoblue (Life Tech). Samples were incubated overnight at -20°C prior to centrifugation. RNA pellets were washed in 75% ethanol and air-dried on ice for 10min before resuspending in RNAse-free water. RNA was treated with DNAseI (Qiagen) to remove any genomic DNA, after which the RNA was cleaned up and concentrated over a RNeasy column (Qiagen).

A low-input method of RNAseq was done based on the Smart-Seq2 method with minor modifications (*72, 73*). In brief, 1 µL of extracted RNA was used for reverse transcription and library preparation. Reverse transcription was done by mixing RNA with 1uL anchored oligo-dT primers and 1uL dNTP (NEB), and denaturing at 72°C for 3 minutes, after which the first-strand reaction mixture was added and incubated in a thermal cycler. PCR preamplication was done by adding the ISPCR primers and the KAPA HiFi HotStart ReadyMix (KAPPA Biosystems) and incubating for a varying number of PCR cycles depending on the sample. PCR product of the cDNA was purified using AMPure XP beads (Beckman Coulter Life Sciences). cDNA quality was checked using a high-sensitivity DNA chip (Agilent) on a Bioanalyzer (Agilent), to measure concentrations and determine median sizes.

cDNA sequencing was done by BMKGENE (Germany) by collecting 20 million 150 bp paired-end sequences by Illumina sequencing. Read quality was assessed using FastQC (www.bioinformatics.babraham.ac.uk/projects/fastqc) and FastP (v1.0.1.; https://github.com/OpenGene/fastp) was used for removing sequencing adapters and low quality reads. Next, using salmon (v1.10.2), we built a transcriptome index based on the latest Ceratopteris reference genome (v2.1, https://phytozome-next.jgi.doe.gov/info/Crichardii_v2_1), which was used for quantification with the trimmed reads.

The RNAseq raw reads have been deposited in the NCBI Short Read Archive (SRA) under the BioProjectID: PRJNA1293105.

**Transcriptome analysis - RNA data analysis**

Heatmaps of expression patterns of developmental regulators was made by initially drafting a list of known *Arabidopsis* regulators ((*26, 28*)), from which orthologous genes in Ceratoperis were identified using Orthofinder (*74*). Normalized expression values from DESEQ2 (*75*) were averaged for developmental stages and plotted.

Dynamic time warp analysis (DTW) and temporal phased gene expression profiling was done as described previously (*26*). Gene pairs were identified using a best blast-hit approach using the R package dtw v1.22-3 (https://www.jstatsoft.org/article/view/v031i07). DTW analysis was performed using the R package dtw v1.22-3. The best blast-hit approach was performed using BLASTP."For Figure S6, *WOX* orthologues were identified based on a constructed phylogeny. Firstly, genes were identified containing the Homeobox domain (PF00046) using Pfam, followed by a BLASTP search against the *Arabidopsis* protein database to retain genes whose best hits correspond to known *AtWOX* members. Multiple sequence alignment was then performed using Clustal Omega, and a phylogenetic tree was constructed using IQ-TREE. As there was no clear one-to-one orthologous relationship, DTW analysis between gene expression profiles was done and combined into a single plot. Code for DTW analysis is available on GitHub (https://github.com/TickingClock1992/DTW-analysis-for-plant-embryogenesis)

**Generation of transgenic GH3::GUS lines**

The *GH3::GUS* auxin signaling-responsive cassette (*76*) was cloned from the pUC19 vector into pGEM-T-Easy cloning vector (Promega Corporation, Madison WI, USA) as an *Eco*RI-*Eco*RI restriction fragment. *GH3::GUS* was subsequently cloned into the Ceratopteris transformation vector pBOMBER (*77*) as a *Not*I-*Not*I fragment (Fig. S10A). *GH3::GUS*-pBOMBER was transformed into *Ceratopteris richardii* strain Hn-n using microparticle bombardment, following the protocol and conditions in Plackett *et al.* (2015). Transgenic sporophytes were selected by screening for antibiotic resistance to Hygromycin B (40 µg/ml).

Transgenic lines were assessed in the $T_1$ sporophyte generation for T-DNA copy number by DNA-blot analysis of the Hygromycin selection marker (HygR) and GUS coding sequence, and for the presence of a full-length *GH3::GUS* cassette by genotyping PCR (Fig. S10B-G). Genomic DNA extraction and DNA-blot analysis was performed as previously described (*60*), using probes previously established for both target cassettes (*60*). All primer sequences are given in Supplemental Table S1.

#### GH3::GUS *auxin response assay*

Responsiveness of *GH3::GUS* to auxin in *Ceratopteris* was tested in the $T_2$ sporophyte generation. Transgenic sporophytes were identified within each line by selecting 10 day-old sporophytes under 40 µg/ml hygromycin selection. At 20 days old hygromycin-resistant sporophytes were transferred to sterile C-fern liquid medium (pH6.0) containing mock treatment (0.001N NaOH), 50 µM 1-Naphthaleneacetic acid (NAA) (0.001N NaOH) or 50 µM N-1-naphthylphthalamic acid (NPA) (0.1% DMSO v/v). Gentle vacuum was applied twice for one minute, after which sporophytes were incubated under standard growth conditions for 40 hours prior to GUS staining.

#### *GUS staining*

GUS activity was analysed in the $T_2$ generation. In all GUS staining experiments a *Ceratopteris 35S::GUS*-pBOMBER transgenic line (*77*) was included as a positive control. Sporophyte tissues were fixed in ice-cold 90% acetone for 10 minutes, washed and then stained using 1mg/ml (w/v) 5-bromo-4-chloro-3-indolyl-b-D-glucuronic acid (Melford, Ipswich, UK) at 37°C for 16 hours. Reagent uptake was encouraged by applying gentle vacuum twice for five minutes prior to incubation. To capture GUS expression during embryo development populations of 12 day-old whole gametophytes were GUS-stained at daily intervals between 0 and 7 days after flooding with sterile water to induce fertilization. Gametophytes were not fixed with acetone and stained at 37°C for 16 hours with 1 mg/ml (w/v) 5-bromo-4-chloro-3-indolyl-b-D-glucuronic acid and 5 µM potassium ferricyanide to restrict stain migration, applying gentle vacuum twice for four minutes prior to incubation. GUS-stained sporophyte and gametophyte tissues were decolourized by repeated incubation in 70% (v/v) ethanol at room temperature. Sporophytes were dried and imaged under a dissecting stereomicroscope, gametophytes were rehydrated and imaged with a Zeiss Axioplan microscope (Carl Zeiss Microscopy Ltd., Cambridge, UK), both mounted with QImaging MicroPublisher 3.3 RTV cameras (Teledyne QImaging, Surrey, BV, Canada). Images were minimally processed in Photoshop 2024 (Adobe, San Jose, CA, USA) to adjust brightness and contrast.

#### Finite-element modelling of tissue-wide stresses - Calculations of mechanical stress distribution

We use the finite element method to calculate the distribution of mechanical stress in the plant tissue for given spatial variations in cell wall stiffness and a spatially constant turgor pressure. In our calculations, we account for both the vertical cell walls (the walls perpendicular to the plane of the tissue that form the boundaries between neighbouring cells), and the horizontal cell walls (the in-plane walls on the top and bottom of the tissue). As starting point for the modelling, a whole gametophyte was used which cell walls were imaged using SR2200. A max projection of a Z-stack was used to manually trace the cell wall network in Adobe Illustrator (https://www.adobe.com/products/illustrator.html). The images were binarized for further analysis.

**Finite-element modelling of tissue-wide stresses - Image skeletonization and network reconstruction**

First the cellular network is extracted from microscopy images using a custom MATLAB script, which converts a binarized image of the plant tissue into a structured representation of nodes, edges (representing the vertical cell walls), and polygonal cells. A watershed segmentation is applied to delineate cell regions, after which cell wall contours are extracted by skeletonizing the segmented image using morphological thinning. This process reduces the wall structures to single-pixel-wide lines while preserving their topology. Next, branch points (locations where three or more walls meet) are identified using morphological analysis and associated with adjacent cells using dilation and querying the cell labels, so that each cell is linked to a set of branch points that define its boundary. Cell boundaries are then traced on the skeletonized image and intermediate nodes are inserted at regular intervals, so that each cell wall is represented by a set of connected nodes. Boundaries of the tissue are identified as those cell walls that are associated with only one cell. Finally, each cell in the tissue is triangulated using the DistMesh2D function (*78*) to generate a background mesh that represents the horizontal cell walls.

**Finite-element modelling of tissue-wide stresses - Finite element calculation**

We performed finite element calculations to calculate the patterns of mechanical stress in the tissue resulting from the mechanical equilibrium between the turgor pressure inside the cells and the cell wall elasticity. We assumed a constant turgor pressure $P$ throughout the tissue, so that the effect of the turgor can be accounted for by an effective imposed normal stress $-P$ on the boundaries of the tissue. Assuming that the vertical cell walls are homogeneous across the thickness of the tissue, each segment of a vertical cell wall (discretized as described above) is modelled as a linearly elastic mechanical bar element with a stiffness $k_i = E_i hd/\ell_i$, where $E_i$ is the Young's modulus of the cell wall, $h$ the cell wall thickness, $d$ the thickness of the tissue, and $\ell_i$ the length of segment $i$. The horizontal cell walls are modelled as a linearly elastic isotropic material with Young's modulus $E$, Poisson ratio $v = 0.3$ (*79*) and thickness $h$. The relative importance of the vertical and horizontal cell walls for determining the deformation field of the tissue depends on the relative stiffnesses and dimensions of the two types of walls, as expressed by the ratio $\alpha = E_V d h_V / 2 E_H h_H^2$, with $E_V$ and $E_H$ the Young's moduli of vertical and horizontal cell walls, respectively and $h_V$ and $h_H$ the corresponding cell wall thicknesses. We show calculations for $\alpha = 60$ (Fig. S21), but have verified that the results are qualitatively similar for different values of $\alpha$. We use our calculations to investigate how gradients in cell wall stiffness influence the pattern of mechanical stress. We assume that the stiffness depends on the distance to the notch $r$ according to

$$E(r) = E_\infty + \Delta E e^{-r/\xi} \tag{1}$$

where $E_\infty$ is the Young's modulus of the mature tissue, far away from the notch and $\Delta E$ the difference in modulus of the cell walls at the notch. For $\Delta E > 0$ the cell walls close to the notch are stiffer than the mature cell walls, while for $\Delta E < 0$ the cell walls near the notch are softer. The decay length $\xi$ sets the length scale over which the stiffness varies.

We then minimize the total energy to obtain the local strain and stress in mechanical equilibrium. The local stress tensor $\boldsymbol{\sigma}(\boldsymbol{r})$ is diagonalized to obtain the principal stresses $\sigma_1$ and $\sigma_2$, corresponding to the maximum and minimum tension at each location. The direction of the maximum tension is specified as the angle $\theta$ between the direction of the first principal stress and the direction towards the notch. The local anisotropy of the stress is calculated as $(\sigma_1 - \sigma_2)/\overline{\sigma}$, where $\overline{\sigma} = (\sigma_1 + \sigma_1)/2$ is the average stress. Note that within the framework of linear elasticity, all stresses are proportional to the the magnitude of the turgor pressure and both the principal directions and the stress anisotropy do not depend on the value of $E$ and $P$.

**Chemical treatments - Exogenous auxin treatments of embryos**
Auxin treatments were done in two different ways. Initially, gametophytes were transferred to ½ MS plates supplemented with exogenous auxin and subsequently flooded with liquid ½ MS (±10 ml) supplemented equally with exogenous auxin. To ensure proper entrance of exogenous auxin. Later experiments were repeated by again transferring sexually mature gametophytes to ½ MS plates supplemented with exogenous auxin and left for 6-8 hours, before fertilizing with liquid ½ MS supplemented with exogenous auxin. For auxin treatments, the natural indole-acetic acid (IAA, Alfa aeser) was used or the synthetic auxin 1-naphthaleneacetic acid (NAA, Sigma). Additionally, the auxin transport inhibitor N-1-naphthylphthalamic acid (NPA, Sigma) was used. All chemicals were dissolved in a DMSO stock. Embryos were collected by harvesting the gametophytes after one, two or three days after fertilization and cleared and imaged as described previously.

**Chemical treatments - NAA and NPA treatments for altered morphology**
Spores were sown on ½ MS medium supplemented with exogenous NAA (5µM) or NPA (5µM) and grown for approximately twelve days (***50***). For modeling, plants were cleared as described previously and their cell wall structure was imaged and segmented. For embryo orientation, gametophytes were moved from auxin-plates to normal plates and fertilized with water. After two days gametophytes were collected, cleared and imaged for embryo orientation.

**Surgical ablations**
Gametophytes were moved to a sterile petri dish under a Motic stereoscope in a laminar flow hood. Microlances (BD) were used to make incisions/ablations on either the basal, notch, or on the lateral wings next to the region of archegonia of the gametophytes. Attention was paid to not damage the gametophytes more than necessary. In practice, we damaged approximately 5-10 cells. After ablations, plants were moved to a new plate of ½ MS and directly flooded with liquid ½ MS. Control plants were directly moved from the original plate to the new plate. After one or two days, plants were harvested, cleared and subsequently imaged and quantified for embryo development.

**Data visualization**
All data were plotted and visualized in R using the ggplot2 package (https://cran.r-project.org/web/packages/ggplot2/index.html), except the Heatmap which was plotted using ComplexHeatmap (Bioconductor - ComplexHeatmap).

**Supplementary Figures**

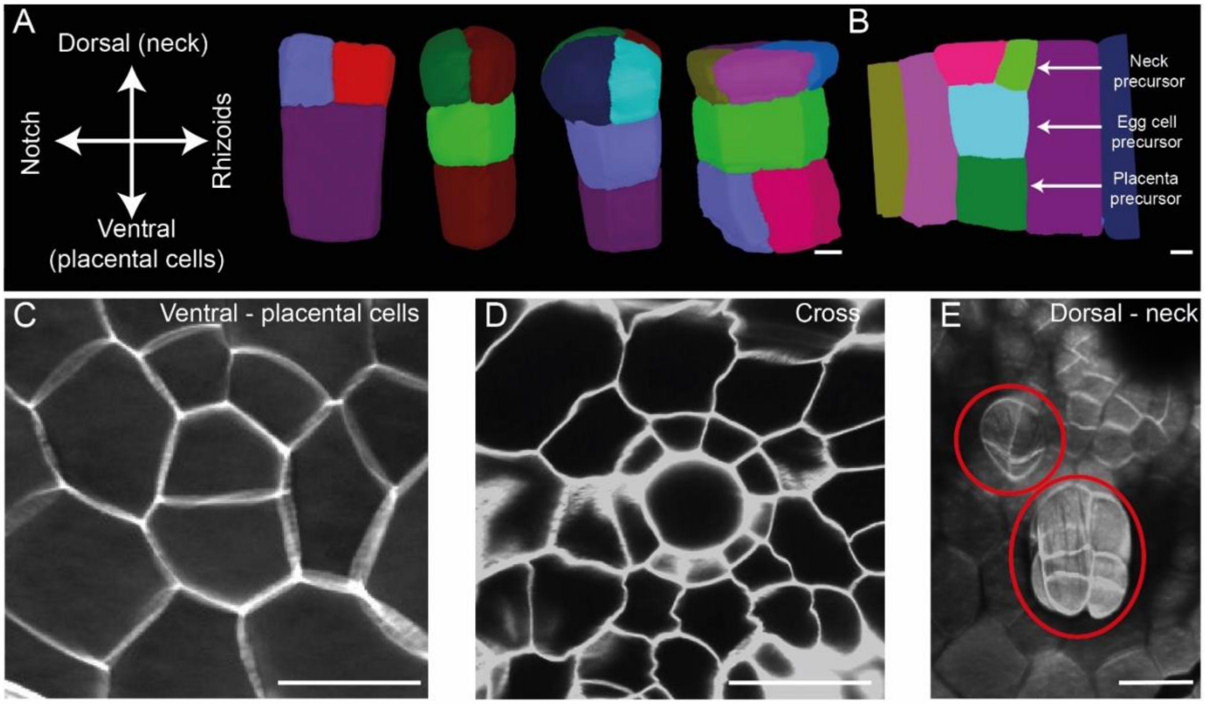


**Fig. S1. 3D overview of archegonium development and structure.**
**(A)** Developmental time series (lateral/side view) of archegonium development: a single cell divides along its long axis to establish the neck precursors on top, the egg cell in the center, and placental cells at the bottom. (**B**) A developing archegonium within the maternal tissue. (**C-E**) In-plane sections through maternal tissue at the level of the placental cells (**C**), the egg cell (**D**) and the necks of mature archegonia (**E**). Scale bars are in (A) 5 µm, (B) 5 µm, (C) 30 µm, (D) 30 µm and (E) 30 µm.

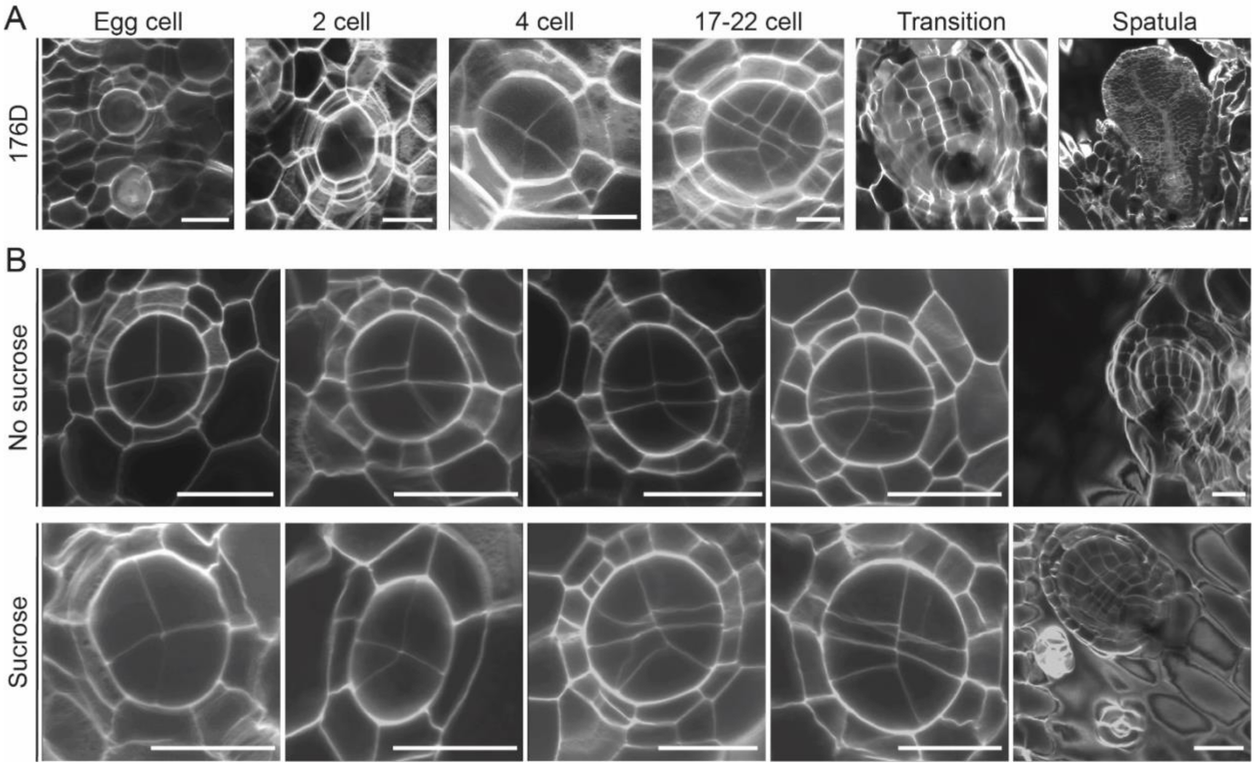


**Fig. S2. Embryo morphology in a second *Ceratopteris* ecotype and Hn-n ecotype under varying growth conditions.**

**(A)** Different embryonic stages in the *Ceratopteris* 176D ecotype. (**B**) Comparison of embryos (Hn-n) developing on gametophytes grown on ½ MS without (top) or with (bottom) 1% sucrose added. Sucrose is known to induce apogamy if gametophytes are grown for longer time periods on it. Scale bars are in (A) 30 µm and (B) 50 µm.

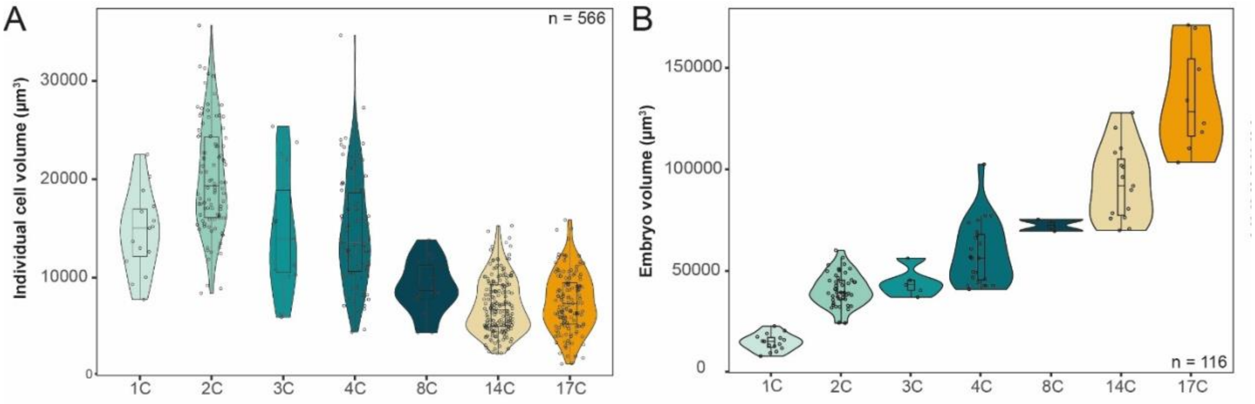


**Fig. S3. Growth of embryos and their cells.**

**(A)** Volume of individual cells from different time points (x-axis shows the number of cells in each embryo) (**B**) Volumes of total embryos from different developmental time points.

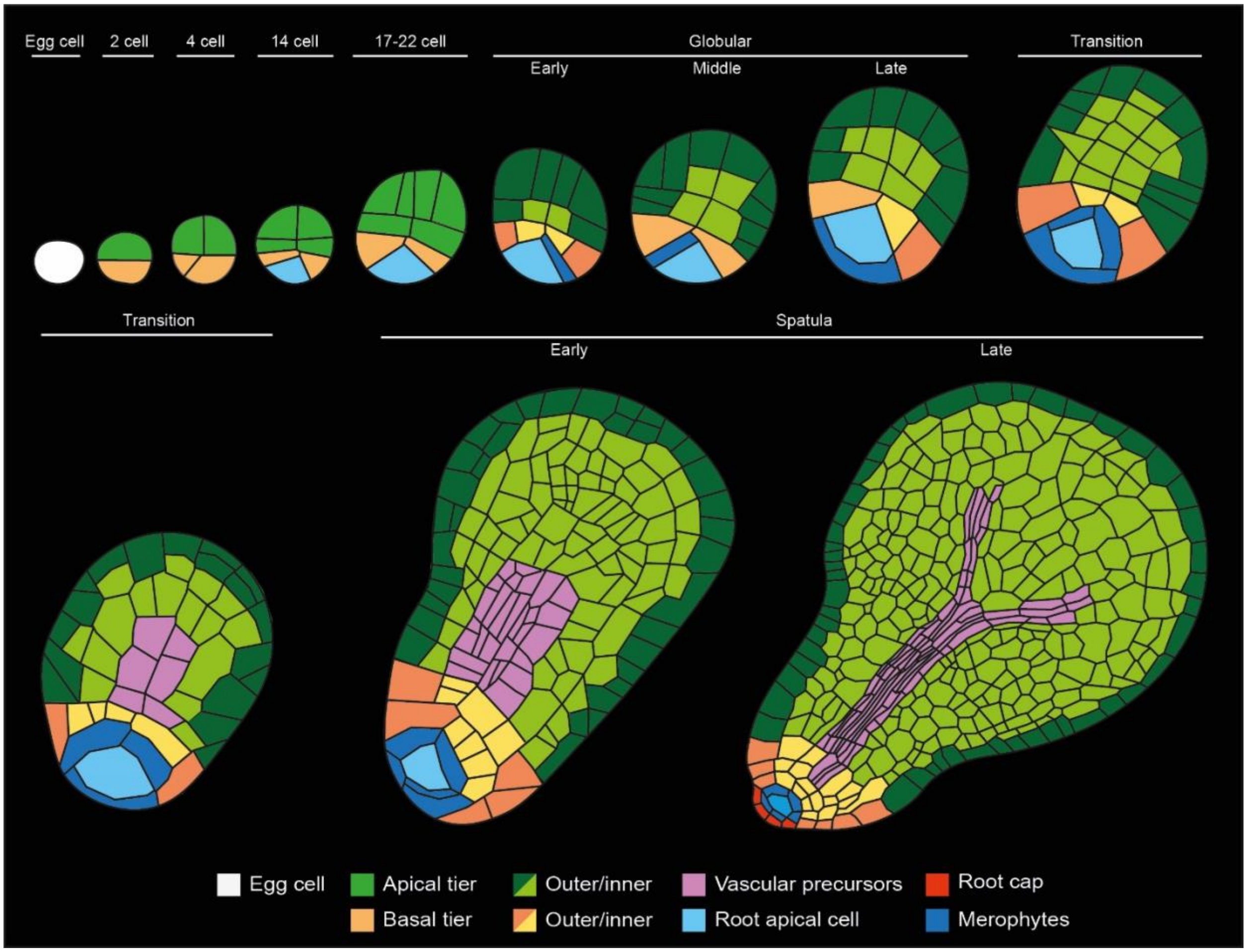


**Fig. S4. Inferred cell lineages during embryo development and organ specification.** Illustrated cross-sections of different embryonic stages in *Ceratopteris*. The different cell lineages and major organs they differentiate to are color coded.

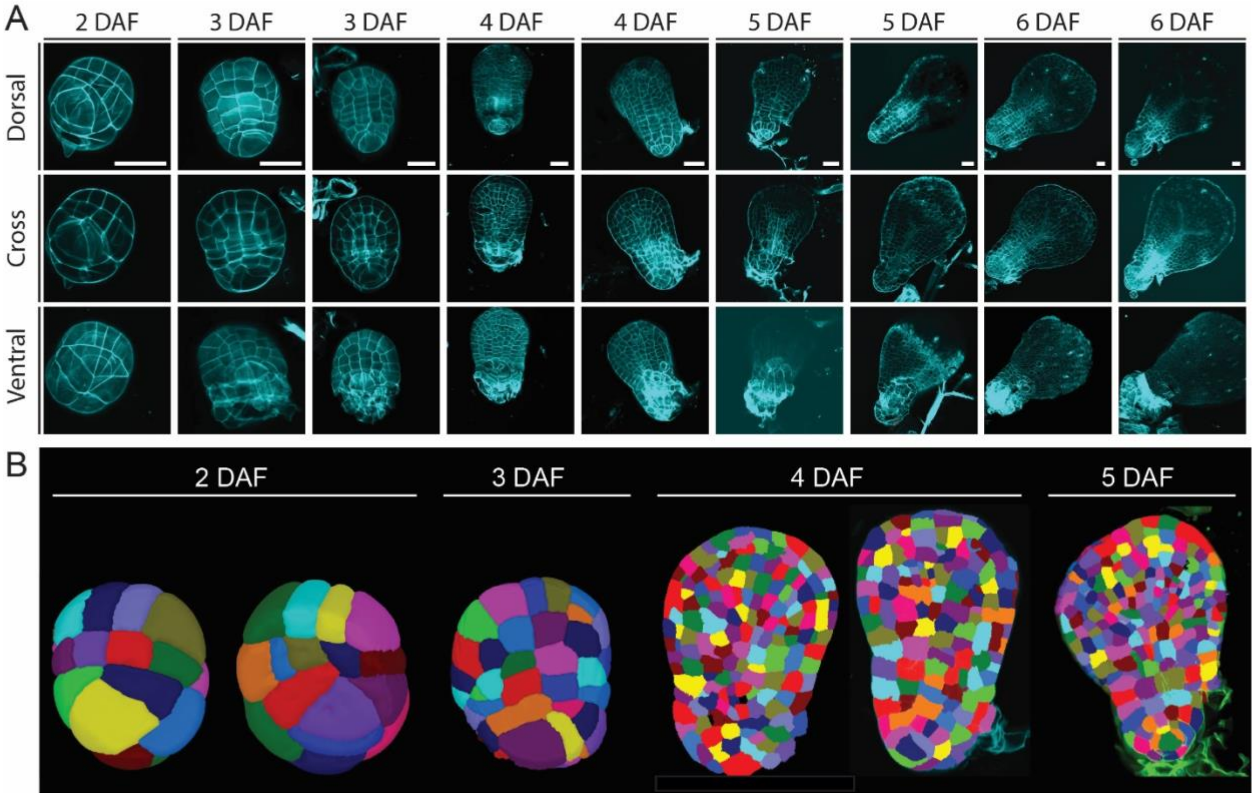


**Fig. S5. Embryonic stages collected for RNA-seq.**
(**A**) Confocal z-slices of embryos from the different time points collected for RNA-seq with 3 planes depicting the dorsal, ventral and median cross-section of the embryos and (**B**) 3D segmentation of isolated embryos for two and three days after fertilization (DAF) and cross-sections of four and five DAF. Scale bars in (A) 50 µm.

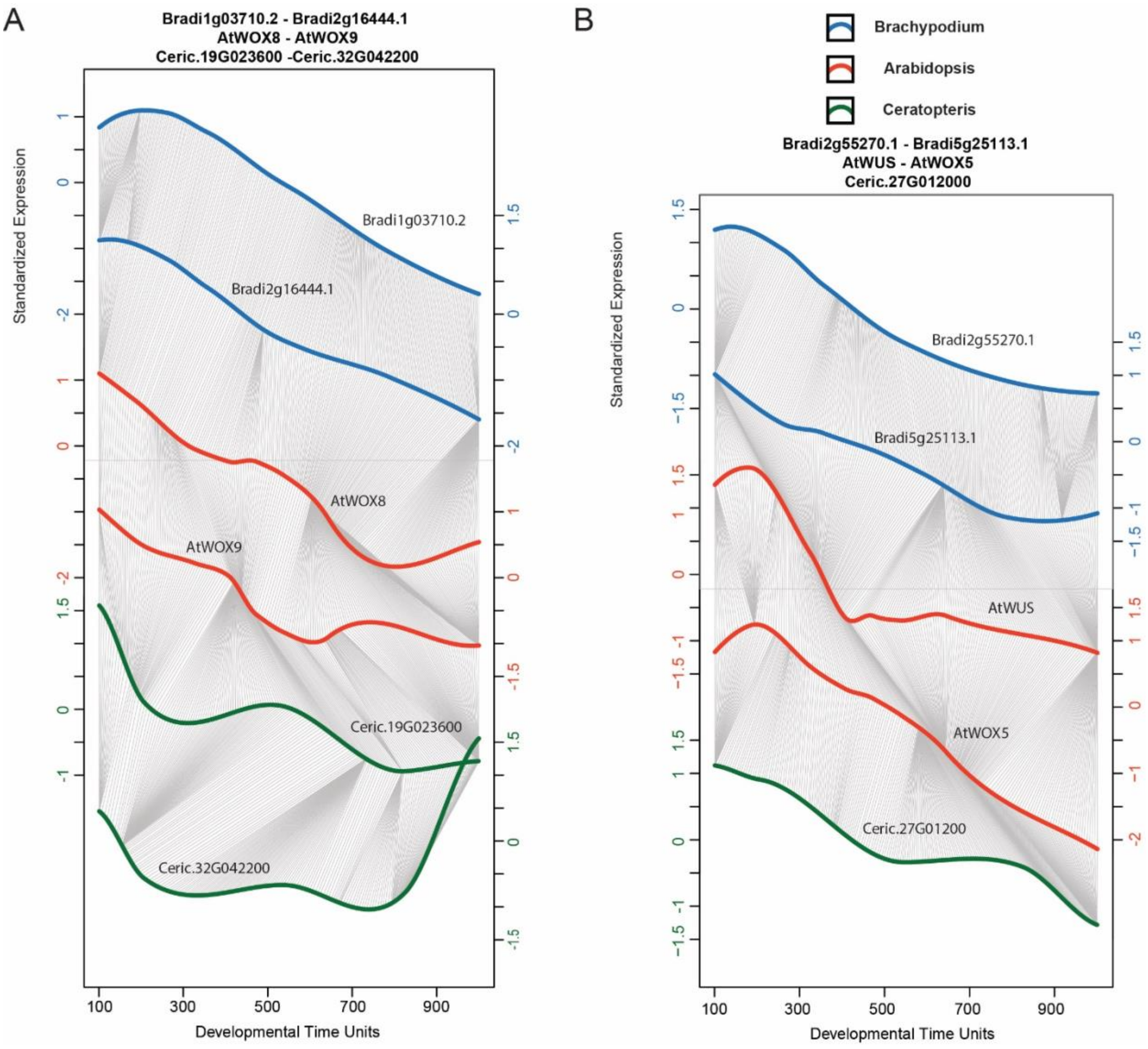


**Fig. S6. Dynamic time warp analysis of *WOX* genes.**

Dynamic time warp expression patterns across embryogenesis of **(A)** *Arabidopsis WOX8* and *WOX9* genes, compared to *Brachypodium* orthologues and two *Ceratopteris* orthologues, and (**B**) *Arabidopsis WUSCHEL* and *WOX5* genes and *Brachypodium* and *Ceratopteris* orthologues.

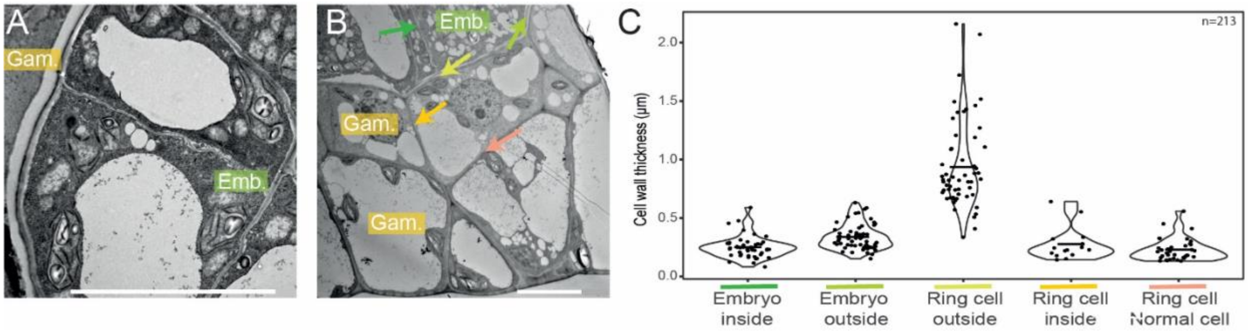


**Fig. S7. Cell wall thickening around the embryo.**
**(A)** TEM image of the embryo and the gametophyte boundary showing a darkly stained layer around the embryo. (B) TEM overview image of the embryo, gametophytic ring-shaped cells around it and gametophytic maternal tissue around that. Color-coded arrows depict different cell walls whose thickness was quantified in (C). (**C**) Quantification of cell wall thickness of specific cell walls. Scale bars in (A) and (B) are 10 µm.

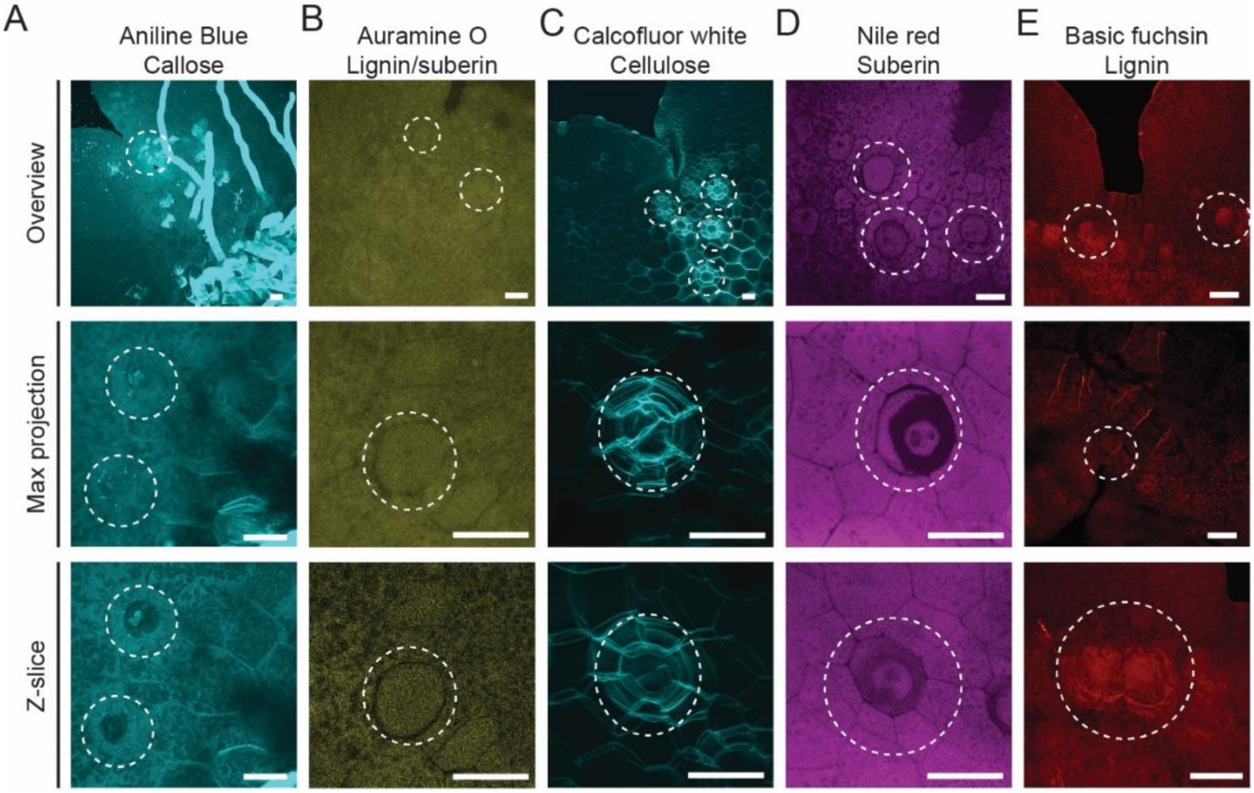


**Fig. S8. Cell wall staining of the archegonium.**

**(A-E)** Staining of cleared mature gametophytes with a focus on the archegonium. Top panel: overview of the notch meristem and archegonia; Middle panel: maximum projection of the archegonium; Bottom panel: cross-section through the egg cell. Archegonia were stained to test for different cell wall polymers with (A) Aniline Blue for Callose, (B) Auramine O staining Lignin/suberin, (C) Calcofluor white for cellulose, (D) Nile red or suberin and in (E) Basic fuchsin staining for lignin. Scale bars in (A), (B), (C), (D) and (E) are 25 µm.

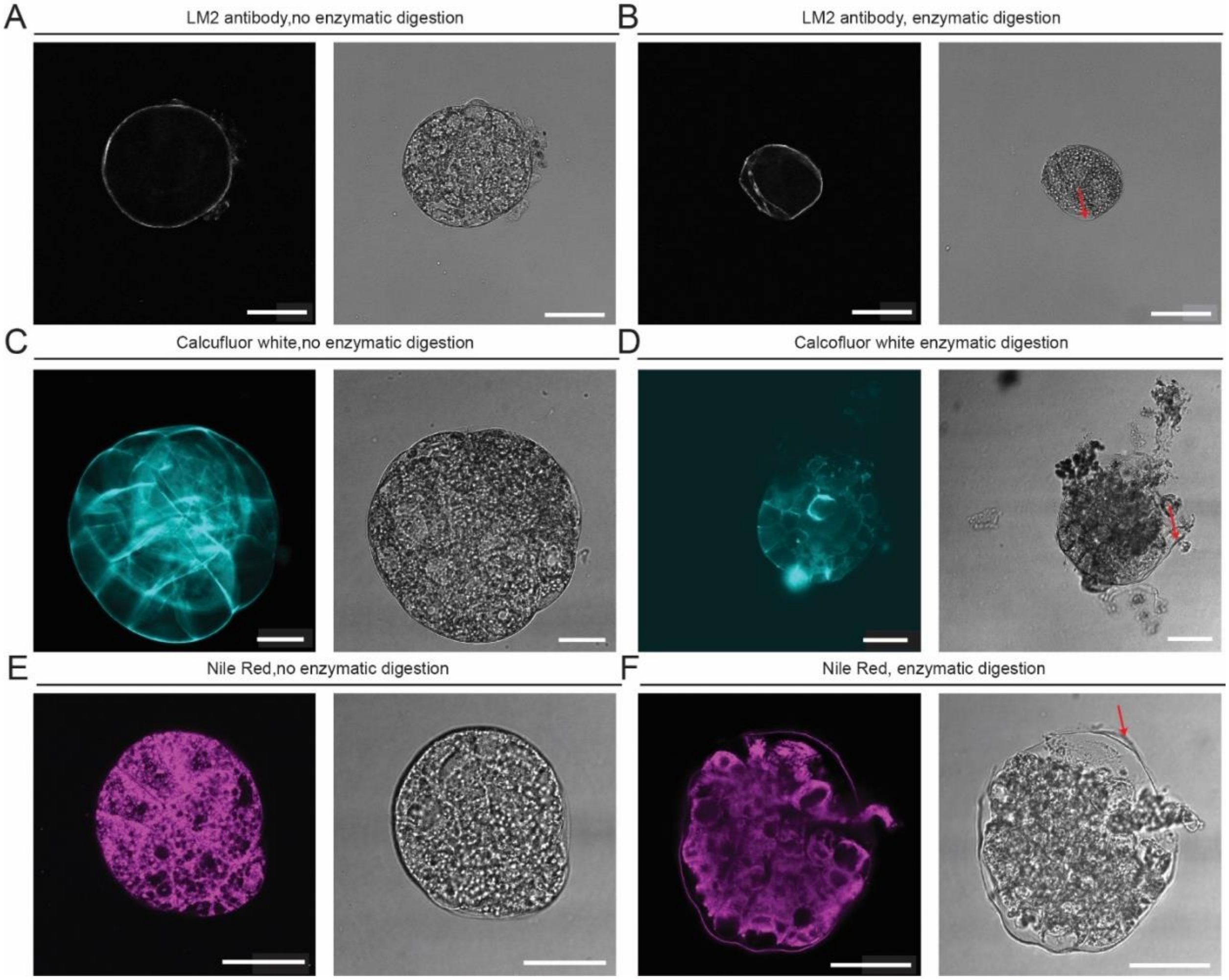


**Fig. S9. An envelope-like structure surrounds the *Ceratopteris* embryo.**
(**A-F**) Staining of cell wall material of undigested isolated embryo's (A,C,E) or enzymatically digested embryos (B,D,F). Left panel shows fluorescence signal, while right panel shows bright field image, with (A,B) LM2 immunostaining for arabinogalactan protein, (C,D) Calcofluor white staining for cellulose and (E,F) Nile Red staining for lipids. Red arrows indicate the envelope-like structure. Scale bars in (A), (B), (C), (D), (E) and (F) are 40 µm.

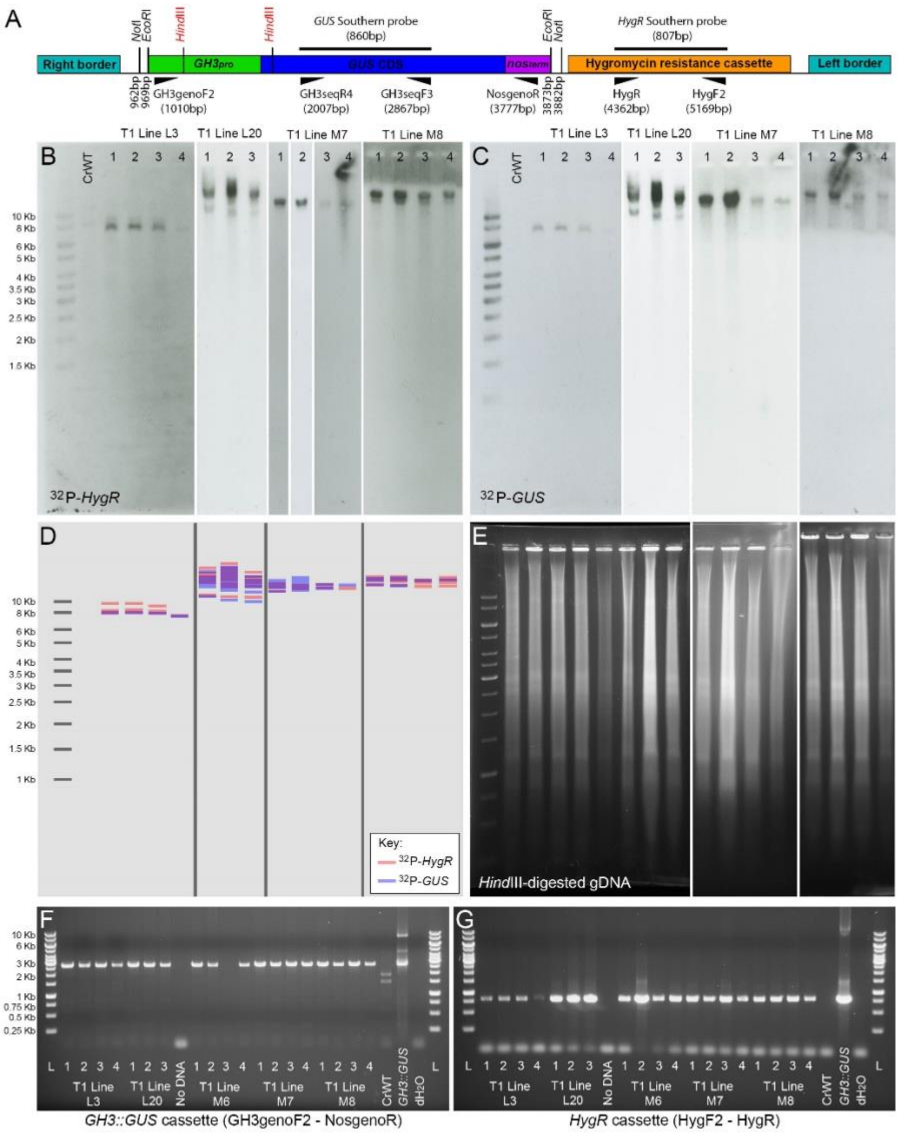


**Fig. S10. Molecular validation of *Ceratopteris* p*GH3::GUS* reporter lines.**

(**A**) Schematic of the pGH3::GUS-pBOMBER T-DNA (6408bp total length), showing positions of restriction sites for cloning and DNA blot analysis, the sites of probe hybridisation (probe sizes in brackets) and genotyping/probe primer binding sites (positions in brackets). Restriction site and primer positions are with reference to the start of the Right Border sequence. (**B-E**) T1 sporophyte DNA gel blot analysis, showing hybridisation patterns of P-32 radiolabelled Hyg and GUS cassette probes (B,C) and their overlap (D, schematic diagram) when probing HindIII-digested genomic DNA from each individual (E). C and D represent repeat probing of the same membrane-bound genomic DNA. All images scaled according to the visible DNA ladder. Four independent T1 lines are shown, with 3-4 sibling sporophytes tested per line. (**F,G**) Genotyping PCR of T1 sporophytes for the full length GH3::GUS cassette (F) and HygR (G) using the primers shown in (A). T1 lines L3, L20, M7 and M8 carry 1-4 copies of the GH3::GUS cassette including at least one full-length. Transgene patterns are similar between siblings of the same T1 line.

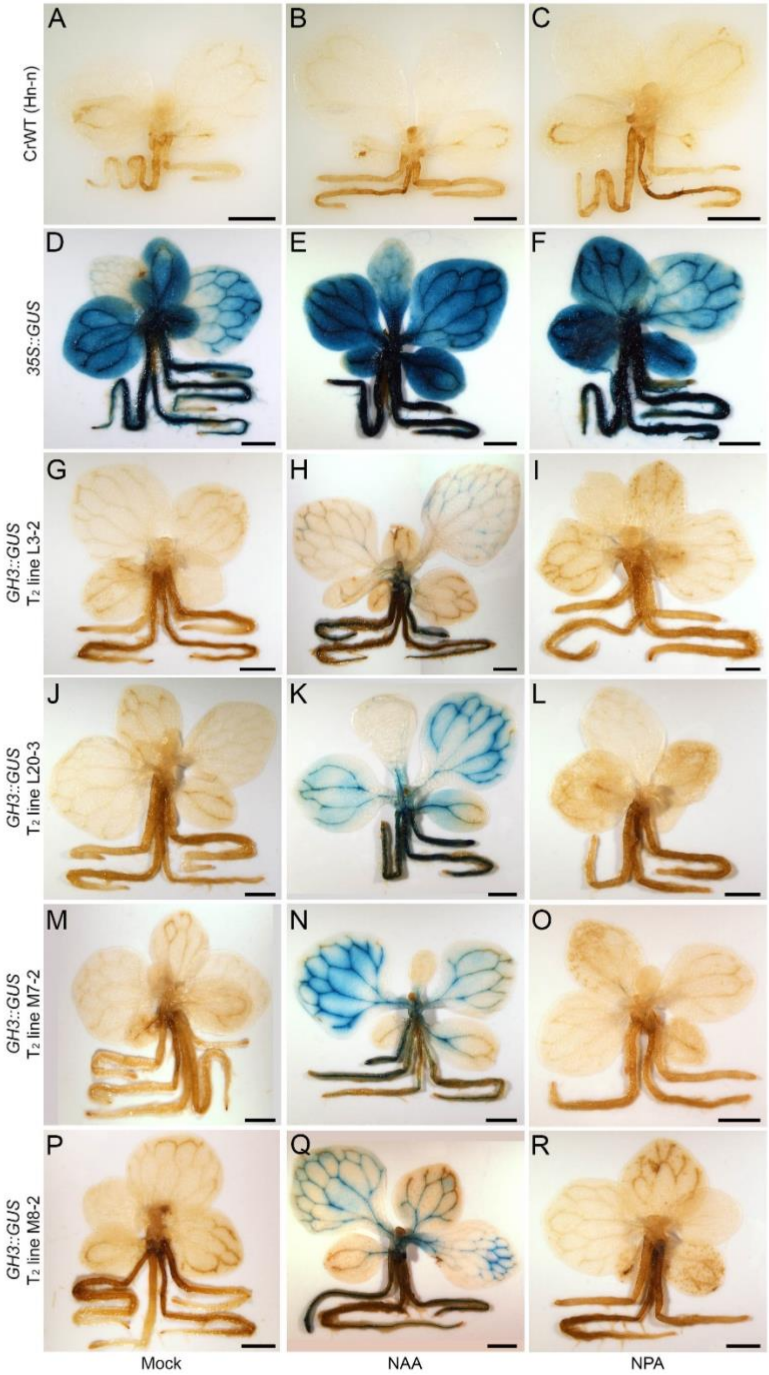


**Fig. S11. Auxin-responsive expression of p*GH3::GUS* in *Ceratopteris* sporophytes.** Whole GUS-stained 20 day-old sporophytes representing wild-type (**A-C**), 35S::GUS (**D-F**) and four GH3::GUS T2 lines (L3-2 [**G-I**], L20-3 [**J-L**], M7-2 [**M-O**] and M8-2 [**P-R**]) descended from validated transgenic T1 individuals (Supplemental Fig. 10). Two days prior to GUS-staining sporophytes were infiltrated with mock treatment (A, D, J, M, P), exogenous auxin (50 μM NAA) (B, E, H, K, N, Q) or exogenous auxin transport inhibitor (50 μM NPA) (C, F, I, L, O, R). Sporophytes of all four GH3::GUS lines exhibited more pervasive GUS staining across a wider range of tissues after exogenous NAA treatment. Scale bars = 2 mm.

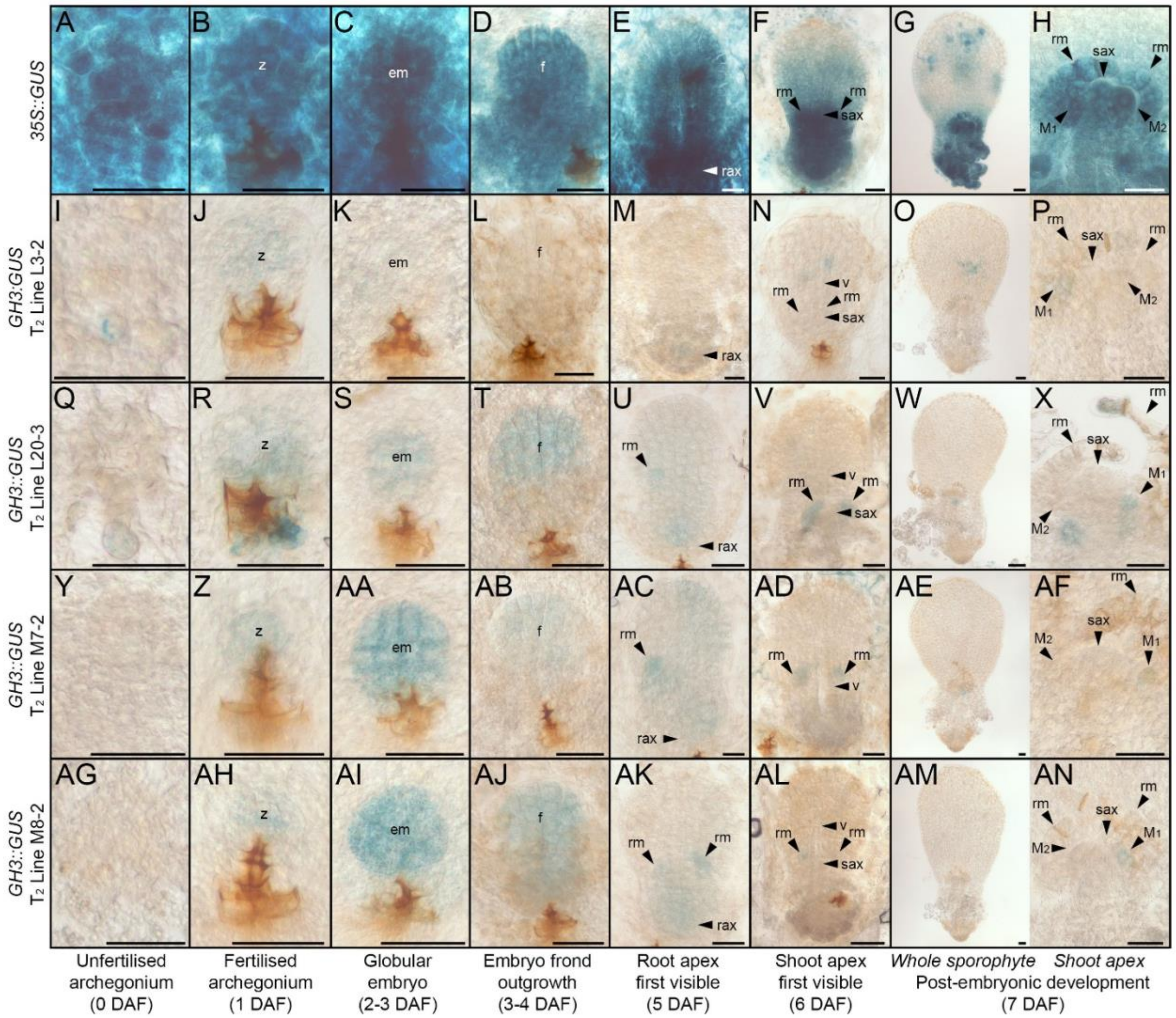


**Fig. S12. *GH3::GUS* expression during *Ceratopteris* embryo development.** GUS staining of *Ceratopteris* embryos through development, comparing staining of 35S::GUS (**A-H**) against four GH3::GUS T2 lines (L3-2 [**I-P**], L20-3 [**Q-X**], M7-2 [**Y-AF**] and M8-2 [**AG-AN**]) descended from validated transgenic T1 individuals (Supplemental Fig. 10). Embryo development within each line is staged by key morphological events (as shown), with approximate ages listed in brackets. Scale bars = 50 µm.

DAF, days after fertilization; em, globular embryo; f, embryo frond; Mn, merophyte (position n); rax, root apex; rm, ramentum (‘scale leaf’); sax, shoot apex; v, vascular strand; z, zygote

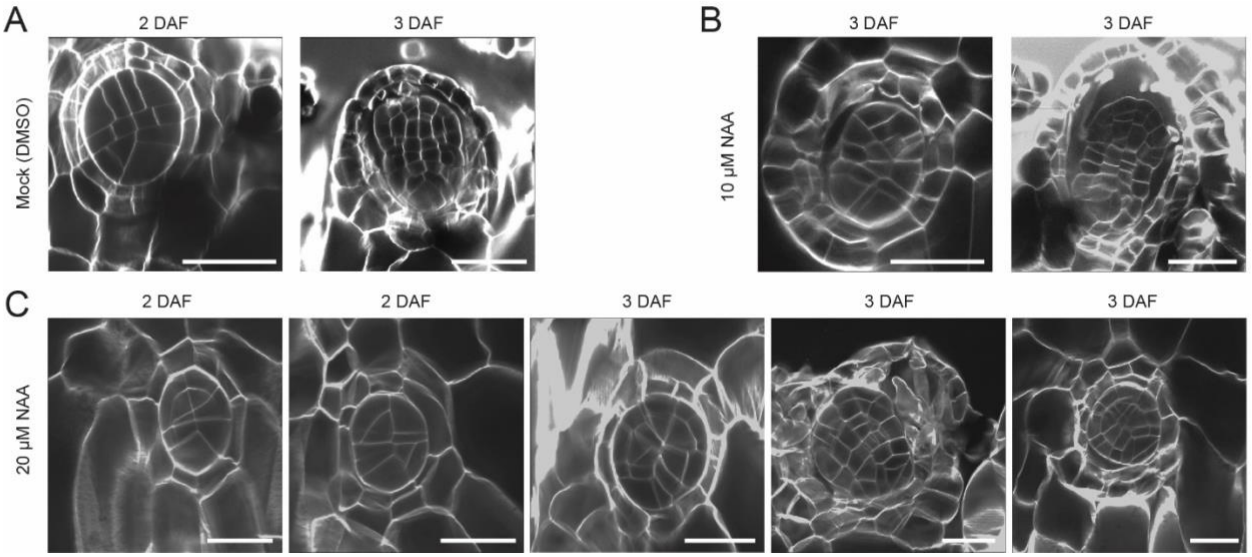


**Fig. S13. Auxin treatment induces abnormal embryo development**
Embryos were imaged two or three days after fertilization (DAF) under different chemical treatments (treatments only started at the moment of fertilization). **(A)** Mock-treated plants. (**B**) 10 µM NAA-treated plants and (**C**) 20 µM NAA-treated plants. Scale bars in (A), (B) are 70 µm and in (C) 50 µm.

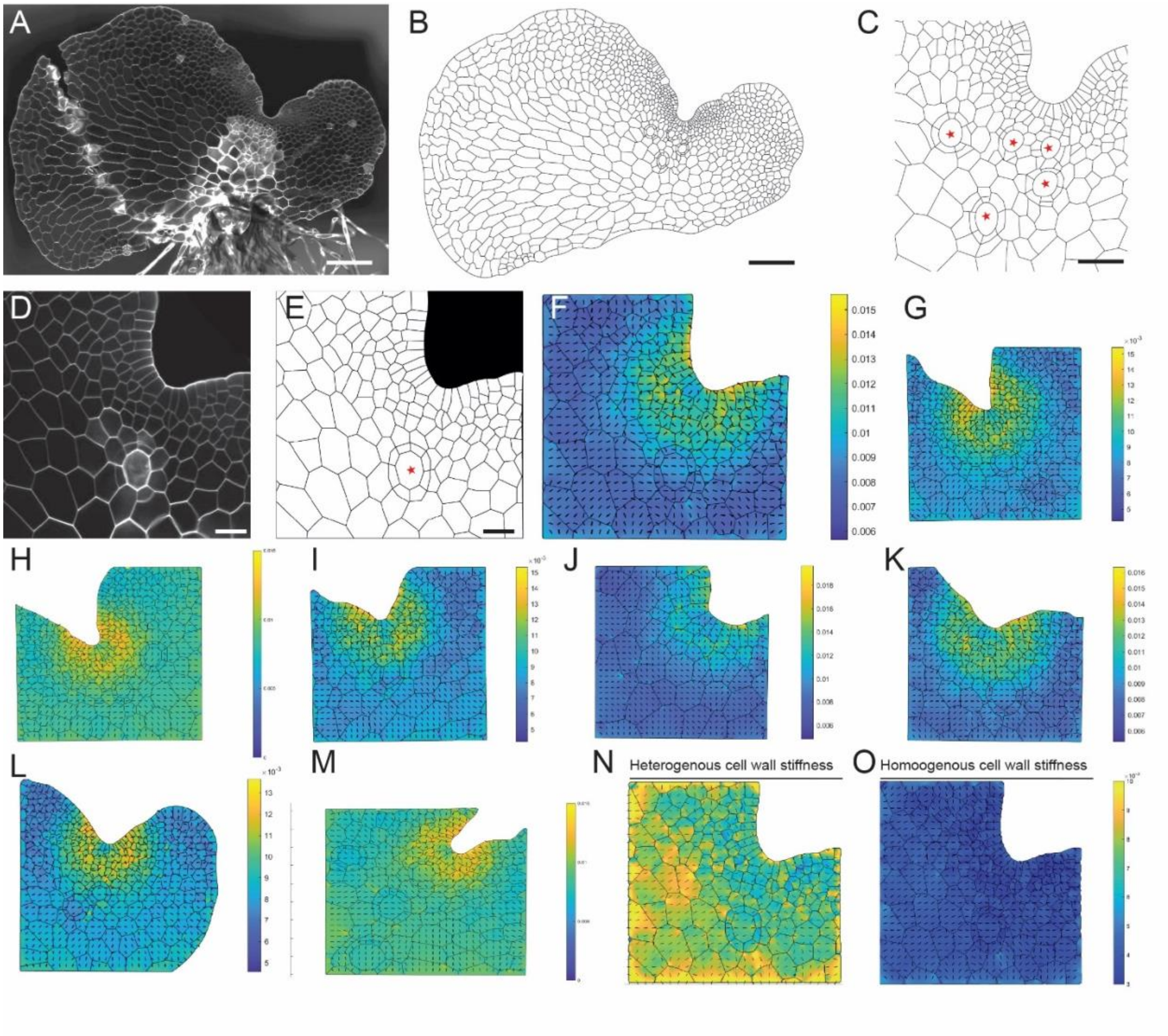


**Fig. S14. FEM modeling of meristematic notches shows a robust stress pattern across individuals.**

(**A**) Maximum projection of whole gametophyte imaged via tile scanning and in (**B**) the extracted cell wall network. (**C**) A close-up of the notch of the plant shown in (B). (**D-F**) A close-up of a different individual and its maximum projection (D) and cell wall network (E) and the modeled stress anisotropy within the notch (F). (**G-M**) Modeled stress anisotropy patterns in seven additional individual plants. **(N-O**) Comparison between stress anisotropy with (N) and without (O) the assumption of a gradient in cell wall stiffness. Scale bars are in (A) and (B) 100 µm, (C) 50 µm, (D) 25 µm and (E) 25 µm. Red stars in (C, E) indicate positions of egg cells within archegonia.

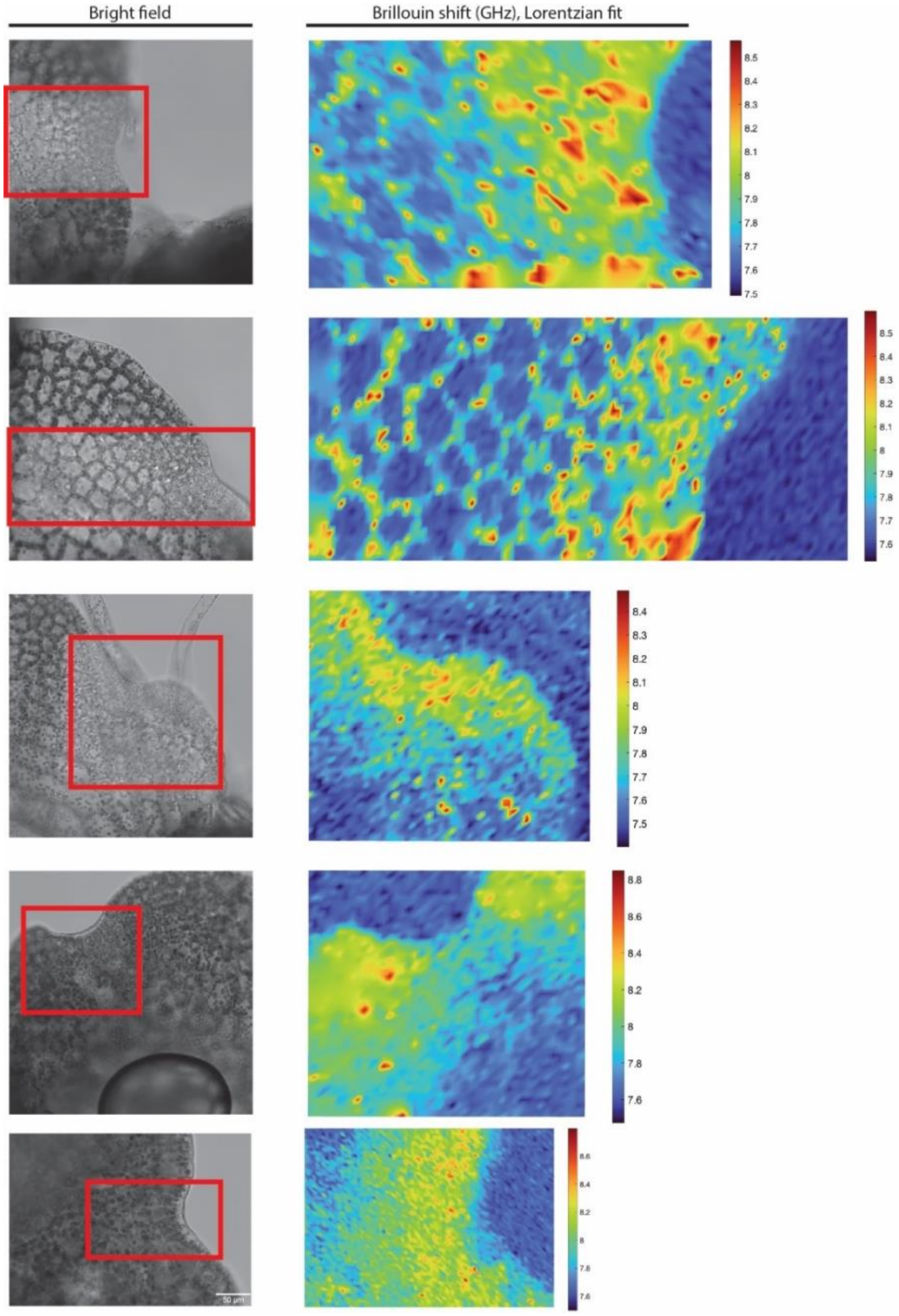


**Fig. S15. A robust Brillouin frequency shift across individuals**
Brillouin microscopy patterns in 5 additional individual plants with brightfield images on the left and on the right the corresponding Brillouin frequency shift (GHz) after a Lorentzian fit. Red boxes depict the area imaged for the Brillouin frequency shift.

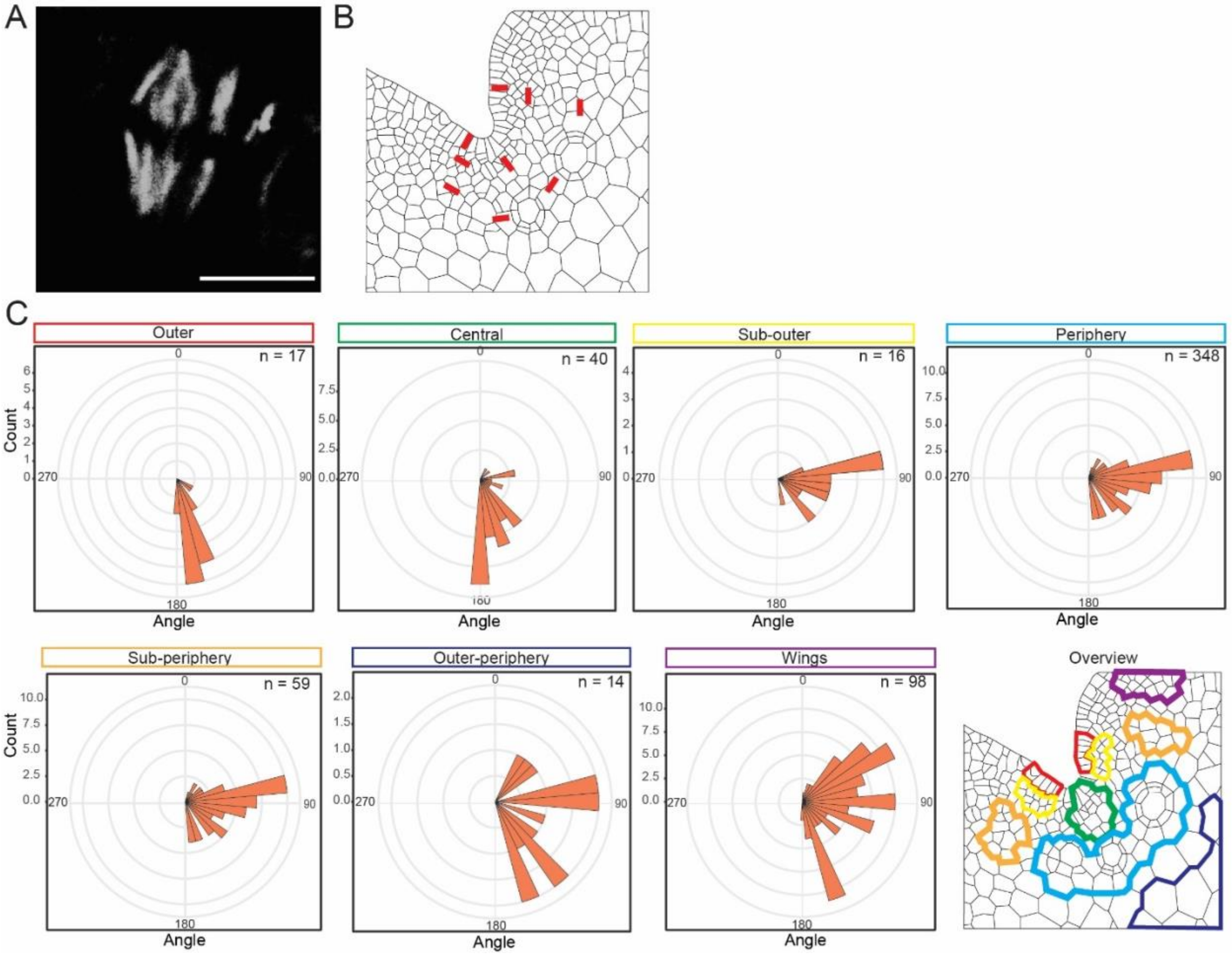


**Fig. S16. Microtubule orientations correlate with predicted stress patterns**
(**A**) Mitotic spindle as a positive control for antibody staining.. (**B**) schematic overview of microtubule orientation throughout the gametophytic notch region. Orientations are based upon (C). (**C**) Quantifications of microtubule orientation in different selected groups of cells. Bottom right panel depicts the different groups of cells and the associated color coding. Scale bars in (A) is 5 μm.

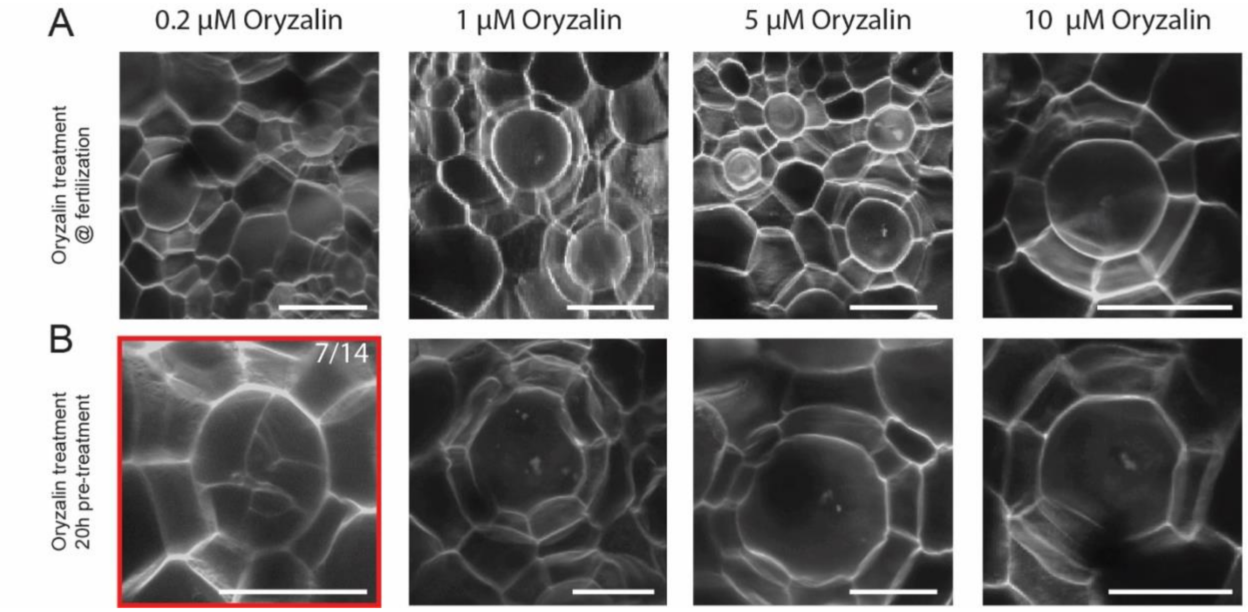


**Fig. S17. Microtubule inhibitors disturb embryo development.**

(**A**) Different concentrations of Oryzalin were added at the moment of fertilization and representative images are shown of embryos at 2 days after fertilization**.** (**B)** Oryzalin pre-treatments were performed by growing gametophytes on medium supplemented with oryzalin for twenty hours, only afterwards plants were fertilized. Oryzalin was not added to the liquid medium used for fertilization, in contrast to (A). Scale bars in (A) are 70 µm and (B) 60 µm. Red box marks the treatment resulting in disrupted embryo development (n=7 out of 14 embryos).

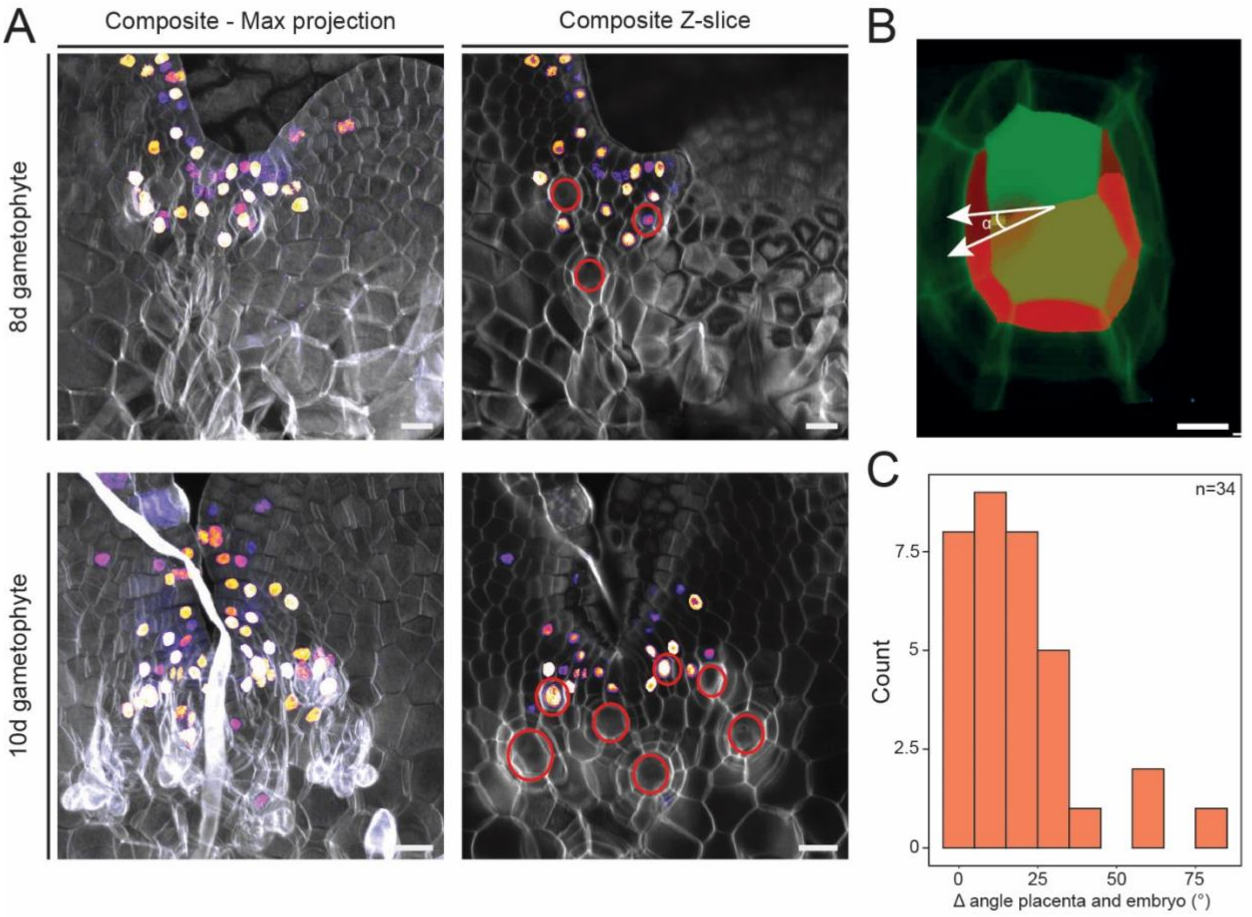


**Fig. S18. Confined cell division patterns in gametophytes show similar orientations as the zygote.**
**(A)** EdU staining of sexually mature gametophytes showing in grey the cell wall staining and purple-orange EdU staining of dividing nuclei**.** (**B**) 3D segmentation of a two-cell embryo and the two placental cells on the ventral side of it, with white arrows depicting the angle between the two cell walls which is quantified in (C). (**C**) Quantification of the angle between the placental cell division and the zygotic cell division with 0° meaning that divisions are parallel. Scale bars in (A) are 25 µm and in (B) 10 µm.

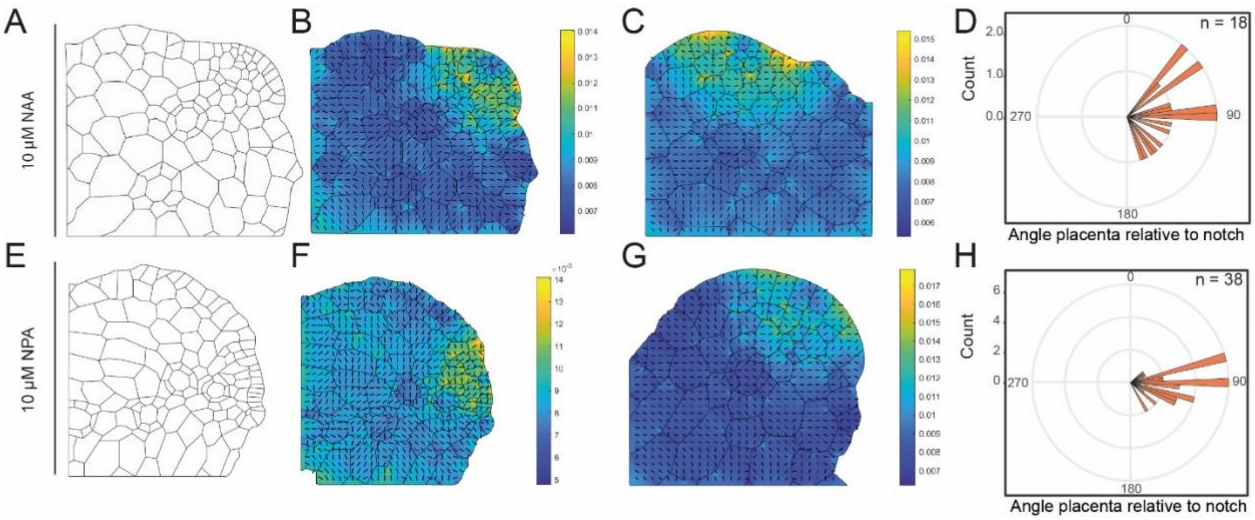


**Fig. S19. Disrupting tissue morphology does not alter the direction of stress anisotropy or archegonium development.**
**(A)** Cell wall network of gametophytes grown on 10 µM NAA with the meristem being the smaller cells on the right top. (**B,C**) Modelled stress anisotropy in two individual plants and (**D**) quantification of the orientation of the placental cell division relative to the notch. This division strongly correlates with the zygotic cell division as shown in Fig S18**. (E)** Cell wall network of gametophytes grown on 10 µM NPA with the meristem comprising the smaller cells on the right top. (**F,G**) Modelled stress anisotropy in two individual plants and (**H**) quantification of orientation of the placental cell division relative to the notch.

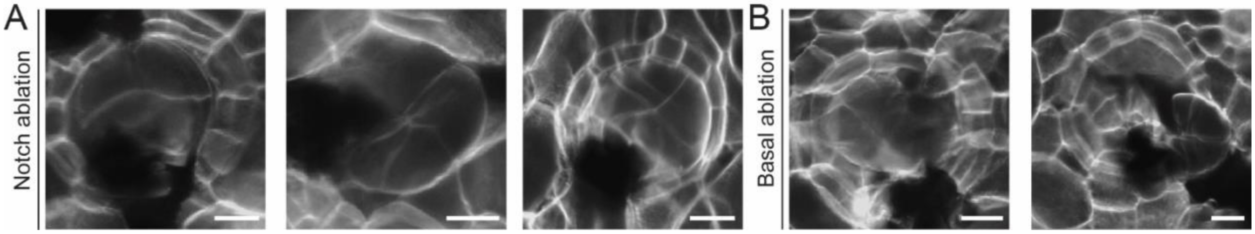


**Fig. S20. Ablating maternal gametophytic tissue can lead to abnormal embryo development (A)** Embryos imaged 2 days after fertilization when the notch was surgically ablated at the moment of fertilization. (B) Embryos imaged 2 days after fertilization when the basal cells (towards the rhizoids) were surgically ablated at the moment of fertilization. Scale bars in (A) and (B) 25 μm

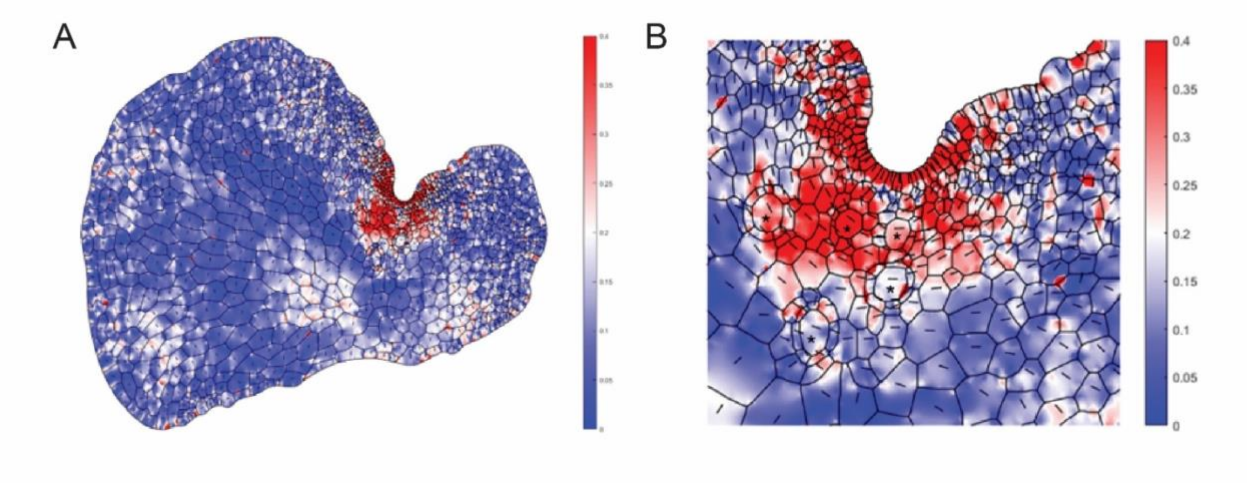


**Fig. S21. Example of finite element calculations and output.**
**(A)** Results of finite element calculations for $\Delta E$=3 and $\xi = 1.2x\ cell\ length$ The line segment in each cell indicates the direction of principal tension, while the color indicates the relative stress anisotropy. (**B**) Zoom in, showing the region close to the notch. The egg cells are indicated with an asterisk.

**Table S1.**
**Primers used in this study**

| Description | Sequence |
|---|---|
| GH3genoF2 | GTAAGAGGGCAAAAATAGGCTG |
| NosgenoR | GGACTCTAATCATAAAAACCCATCTC |
| GH3seqF3 | CTTCGCTGTACAGTTCTTTCG |
| GH3seqR4 | CACTCATTACGGCAAAGTGTG |
| HygF2 | CTTCTACACAGCCATCGGTC |
| HygR | CCGATGGTTTCTACAAAGATCG |

Movie S1.
**Live imaging of *Ceratopteris* embryo from the two-cell stage.**
An embryo grown under microscopy in liquid medium containing 10 µM FM4-64. Observation began at the two-cell stage. Single XY planes were extracted from z-stack images at each time point. The time counter indicates hours and minutes after the start of observation. Scale bar is 20 µm.

**Movie S2.**

**Live imaging of *Ceratopteris* embryo from the two-cell stage.**
An embryo grown under microscopy in liquid medium containing 10 µM FM4-64. Observation began at the two-cell stage. Single XY planes were extracted from z-stack images at each time point. The time counter indicates hours and minutes after the start of observation. Scale bar is 20 µm.

**Data S1.**
Extended data of all raw data used for quantifications plotted in graphs.
